# The fate of mutations on Y chromosomes and autosomes: a unified Wright–Fisher framework accounting for segregation time

**DOI:** 10.64898/2026.04.01.715871

**Authors:** Ariel Offenstadt, Sylvain Billiard, Tatiana Giraud, Amandine Véber, Paul Jay

## Abstract

Understanding how mutations evolve on Y chromosomes is central to explaining the origin, diversity and persistence of sex chromosomes. Mutations occurring on the Y chromosome in sexual populations experience selective dynamics that differ markedly from those on autosomes, due to a reduced effective population size and the presence of large non-recombining regions containing alleles maintained in a permanently heterozygous state. These specific features alter gene transmission in the Y chromosome population compared to autosomes, even within the same pedigree. Here, we provide a two-sex diploid Wright-Fisher model that explicitly incorporates both sex chromosomes and autosomes within a unified population framework, in order to capture the influence of these specificities on the fate of mutations, not only considering fixation probabilities but also segregation times. We use diffusion approximations and provide analytical and numerical tools to compute these quantities across a wide range of parameters and selection regimes. We recover classical results on fixation probabilities in various scenarios, including purely beneficial, deleterious or overdominant mutations, and extend them in the light of mean segregation time, a key but often overlooked determinant of evolutionary outcomes over finite timescales. In particular, our analyses show that overdominant mutations are overall more likely to fix in observable time windows on the Y chromosome than on autosomes. Individual-based simulations corroborate our approximations and highlight parameter regimes where the theoretical approach is particularly useful, especially for parameter values inducing long segregation times or small fixation probabilities, for which simulations are impractical. Our results provide a comprehensive and tractable framework for clarifying how chromosome-specific features shape evolutionary dynamics beyond fixation probabilities alone.

## 1 Introduction

Sex chromosomes play a central role in the evolution of many species. They are frequently involved in speciation processes, contribute to the development and maintenance of sexual dimorphism, and harbor numerous mutations causing sex-linked disorders, such as red–green color blindness in humans. A major explanation for their particular evolutionary impact is that Y (or W) chromosomes are often non-recombining over most of their length and permanently heterozygous. This has important consequences, including the progressive degeneration of the Y (or W) chromosome due to the reduced efficacy of selection in non-recombining regions. Below, we focus on Y chromosomes in XY systems for simplicity, but a similar reasoning holds for W chromosomes in ZW systems (*e*.*g*., in birds). Despite their importance, the consequences of recombination arrest and permanent heterozygosity for the evolutionary trajectories of these regions, as well as for processes such as adaptation, speciation, and genomic conflicts are still poorly known. Similarly, the evolutionary mechanisms driving recombination suppression on Y chromosomes are still debated (Charlesworth, 1978; Saunders and Muyle, 2024; Charlesworth and Olito, 2024; Jay et al., 2025). Moreover, although the rise in genome sequence availability has revealed a striking variation in the genomic architecture, levels of degeneration, patterns of diversity, and evolutionary histories of Y chromosomes across the tree of life, we still lack a clear understanding of the evolutionary forces that have generated and shaped this diversity. For instance, it remains unclear what fraction of the differences in evolutionary trajectories observed between autosomes and sex chromosomes can be attributed to distinct selective regimes affecting these genomic compartments (for example, the occurrence of sex-specific selection), to differences in their recombination landscapes that modulate the strength of selective interference, or to the permanent heterozygosity of the Y chromosome, which influences both the effective population size and the dominance effects of mutations.

These challenges arise in part because we lack a unified framework for rigorously inferring the evolutionary trajectories of mutations on Y chromosomes and for directly comparing these dynamics to those on autosomes, which currently limits our ability to meaningfully interpret the patterns of genetic diversity observed in natural populations. Two key quantities allow a precise description of the fate of a mutation: its fixation or extinction probability and its segregation time, *i*.*e*., the time during which the mutation is neither extinct nor fixed. The latter is often neglected, while it plays a central role in the understanding of the mutation dynamics, most of mutations observed in genomes being still segregating rather than fixed. Moreover, some mutations may have segregation times so long that, even when their fixation probability is substantial, fixation may effectively never occur on observable timescales. For example, strong balancing selection can maintain allele frequencies far from 0 or 1 for extremely long periods, until the stochasticity in reproduction eventually generates a sufficiently large fluctuation to drive the mutation frequency towards one of these absorbing states. This observation raises questions about our ability to directly observe the fixation of new alleles, even in cases where models would predict their fixation with significant probability. Although this issue is well known, it has not been explored in a systematic way to our knowledge.

The fixation probability and segregation time of mutations depend on multiple evolutionary forces, including natural selection and genetic drift. The foundations for understanding fixation and extinction probabilities were established in textbook works by Fisher (1923) and Wright (1931). The Wright-Fisher model assumes that the population size is finite and constant through time and the stochastic nature of the reproduction dynamics is explicitly encoded. This leads to a binary eventual outcome, in which the mutation is either fixed or lost with probability 1. Using a branching process approximation, Haldane proved that the fixation probability of a beneficial autosomal mutation in a Wright-Fisher model could be approximated by a simple quantity equal to twice the selection coefficient (Haldane, 1927). Kimura later extended these results to varying population sizes and strengths of selection, studying the fixation probability as a function of the initial frequency of the mutation (Kimura, 1955, 1962). Although most mutations are known to be deleterious, beneficial mutations have long been considered as key drivers of adaptation to changing environments, and they have been the focus of many models (see Ewens 1967; Patwa and Wahl 2008 and references therein). Deleterious mutations have also been extensively investigated, and shown to accumulate in asexual or selfing populations, for example through Muller’s ratchet (Muller, 1964; Waxman, 2011). They can accumulate in small populations too, even with sexual reproduction, possibly leading to a species extinction (Lynch et al., 1995). In this work, we aim to cover a wide range of selection regimes, including most of the above.

A key element distinguishing the fates of a mutation carried by an autosome and by a non-recombining Y chromosome in a diploid population is the asymmetry between the X and Y chromosomes, present in different proportions in the population. Indeed, in a population with 1:1 sex ratio, only one fourth of the sex chromosomes are Y chromosomes. Although sex chromosomes and autosomes both follow the rules of Mendelian inheritance, a mutation carried by the X or the Y chromosome in the non-recombining region (*i*.*e*., in the sex-linked region) will not be transmitted to males and females in the same way. Consequently, the variations in allele frequencies within these two subpopulations will be different, and will also be different from the variations in allele frequencies experienced by a mutation carried by an autosome. These differences are expected to affect the fixation probability and the segregation time of Y-linked mutations compared to autosomal ones.

By using Haldane’s branching approximations for beneficial mutations and Kimura’s diffusion for deleterious mutations, Charlesworth et al. (1987) compared the substitution rate of these mutations on sex chromosomes and autosomes. They also compared the substitution rate of underdominant mutations on X chromosomes and autosomes. Considering that the substitution rate is equal to the mutation rate times the fixation probability, they found that loci with partially or fully recessive beneficial mutations should show higher substitution rates on Y chromosomes than on autosomes. They also showed that deleterious mutations should fix more frequently on Y chromosomes than on autosomes. However, there has been no such analysis considering overdominant or underdominant mutations on Y chromosomes, *i*.*e*., mutations with an heterozygote advantage or disadvantage compared to both homozygotes. Yet, recent empirical and theoretical works have shown that chromosomal rearrangements, which are abundant on sex chromosomes, may often be overdominant (Berdan et al., 2023). Furthermore, the substitution rate does not account for the differences in segregation times between different mutation types, or between different chromosome types (autosomes versus Y-like sex chromosomes), despite the fact that they condition our ability to actually observe the fixation or extinction of a mutation.

Here, we aim to extend the work of Charlesworth et al. (1987) by deriving expressions for the fixation probabilities and mean segregation times for all types of mutations in autosomes and Y chromosomes. Although several other processes may influence the evolution of sex chromosomes (*e*.*g*., sexually antagonistic selection, Flintham and Mullon 2025), we wish to study the structural effect of the Y-chromosome intrinsic properties, *i*.*e*., its lower effective population size and its forced heterozygosity, on the fixation probability and segregation time of a mutation that it carries. We do not consider mutations appearing on the X chromosome, as the mutation frequency then differs in the male and female subpopulations, and its dynamics in the whole population cannot be satisfactorily described by the same type of one-dimensional Markov processes as those we study here. More precisely, in this case the frequency in the male and female subpopulations must be tracked separately in order to express the law of future changes in mutation frequency, and our methodology based on one-dimensional diffusions does not apply (similar formulae are not available for two-dimensional diffusions).

In order to compare the invasion, fixation and extinction of mutations on autosomes vs. on Y chromosomes, we must consider a diploid, biparental two-sex model. Möhle (1994) introduced a Wright-Fisher biparental two-sex model and studied the probabilities of extinction of the family descending from a given individual at some time in the future. Although one could consider using this result to track the lineage of the original carrier of a particular mutation (and deduce for instance the extinction probability of this mutation), two features make it unsuitable for addressing the question we are interested in with regard to mutation transmission. First, males and females play symmetric roles in this model, preventing the incorporation of key specific features of sex chromosomes. Second, because the genetic transmission tree differs from the biparental pedigree (or family tree, which is the object of interest in Möhle 1994), a family lineage may persist even after a given mutation has been lost. The former limitation also arises in the model proposed in (Möhle and Sagitov, 2003).

To be able to incorporate these specificities in our analysis, we introduce a diploid, biparental, two-sex Wright-Fisher model, in which the two haploid genomes of each individual are picked in exactly one male and one female, accounting for selection and the particular features of heterogametic sex chromosomes. This allows us to finely incorporate the key characteristics distinguishing the allelic transmission for a gene located on the Y chromosome versus an autosome along a biparental pedigree. We derive Wright-Fisher diffusion approximations for the allele frequencies of autosomal and Y-linked genes using the same scaling of time. Indeed, the allele frequencies emerge from this model as discrete-time Markov chains, with complex transition probabilities that render a direct analysis practically impossible, hence the need for diffusion approximations. Using precisely the same timescales in the autosomal and Y-linked contexts allows a direct comparison of the mean segregation times between the two cases. The approach consisting of taking diffusion approximations is the same as Kimura (1962)’s approach and the resulting objects are standard (Durrett, 2008), but we detail how they can be jointly obtained from the same population dynamics. Indeed, the transmission dynamics of both sex chromosomes and autosomes are embedded in the model, building the population pedigree in the same way, and keeping the same selection coefficients, thereby ensuring a legitimate comparison between the fixation probabilities and mean segregating times on the two types of chromosomes. This unified framework also provides numerical and analytical tools to quickly compute these quantities given the selection parameters and the initial frequencies.

The rest of the article is organised as follows. In Section 2.1, we introduce the two-sex biparental Wright-Fisher framework that we will use to model the transmission of an allele carried by an autosome and by the Y chromosome. In Sections 2.2 and 2.3, we describe the procedure we use to derive diffusion approximations for the allele frequency process in the two chromosomal contexts. In Section 3, we analyze and compare the fixation probabilities and mean segregation times derived from the diffusion approximations for a range of parameters including four main selective cases (beneficial, deleterious, underdominant and overdominant mutations). We also show through simulations that the diffusion approximations yield accurate predictions, even for rather small population sizes (*N* = 100 or *N* = 1000), and illustrate the added value of our analytical approach in cases where excessively long segregation times or low fixation probabilities prevent fixation from being actually observed. Finally, our results are discussed in Section 4.

## 2 Methods

### 2.1 Model

We want to compare the probability and time of fixation of mutations under different selection scenarios depending on whether they appear on an autosome or a Y chromosome. To do so, we consider a diploid population of fixed size, composed of *N* individuals with equal number of males and females. The population has an XY sex-determining system, where females are XX and males are XY. The newly arisen mutation, *a*, appears at a particular locus where the ancestral allele (or wild type) *A* is initially fixed. The mutation induces differences in fitness classically encoded as follows:

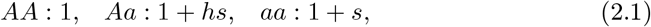

where *h* and *s* are such that *hs* and *s* belong to [−1, 1]. This framework accommodates both a single point mutation and a non-recombining region treated as a single locus and whose selective impact is given by *h* and *s*. We use the same coefficients for a locus carried by the pair of sex chromosomes or by a pair of autosomes.

When we describe the process of gamete formation and fusion underlying our model, we denote the joint inheritance of sex chromosomes and of the alleles at the locus of interest by combinations of two letters, where the first letter is either X or Y depending on the sex chromosome present in the gamete, and the second letter is *A* or *a* depending on the allele also present in the gametic genome. In this notation, when the locus of interest is directly carried by the Y chromosome, gametes can only be of type X*A*, Y*A* and Y*a*. When the locus at which the mutation of interest occurs is carried by an autosome, the association between the sex chromosome and the ancestral or mutant allele is only due to the random formation of gametes and it has no impact on the transmission of the mutation (as we assume that the population sex ratio remains constant through time). In the case of an autosomal locus, gametes can therefore be of type X*A*, X*a*, Y*A* or Y*a* and only the *A/a* state actually matters in the evolutionary dynamics. Nevertheless, we use the same 2-letter notation to describe the genetic content of the gamete in order to have a consistent set of notation in the two scenarios that we want to compare.

The reproduction dynamics is stochastic, with discrete, non-overlapping generations and random mating in the spirit of the classical Wright-Fisher model. In each generation, males and females produce an infinite number of gametes, carrying alleles in the same proportions as those in the current generation. An infinite pool of juveniles is obtained by merging all pairs of paternal and maternal gametes, which are then split into two groups: those who received an X paternal chromosome and those who received a Y paternal chromosome. To form the next generation, *N/*2 males and *N/*2 females are sampled among these juveniles, carrying alleles with probabilities depending on their selection coefficients (see Figure 1). The proportions of the three possible genotypes *AA, Aa* and *aa* in the male juvenile population are denoted by 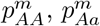 and 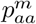, and each male in the next generation will carry genotype *AA, Aa* and *aa*, respectively, with probabilities

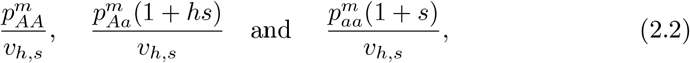

where 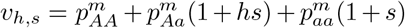, and similarly for females (with genotype frequencies in the female juvenile population denoted by 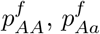 and 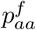).

**FIGURE 1:**
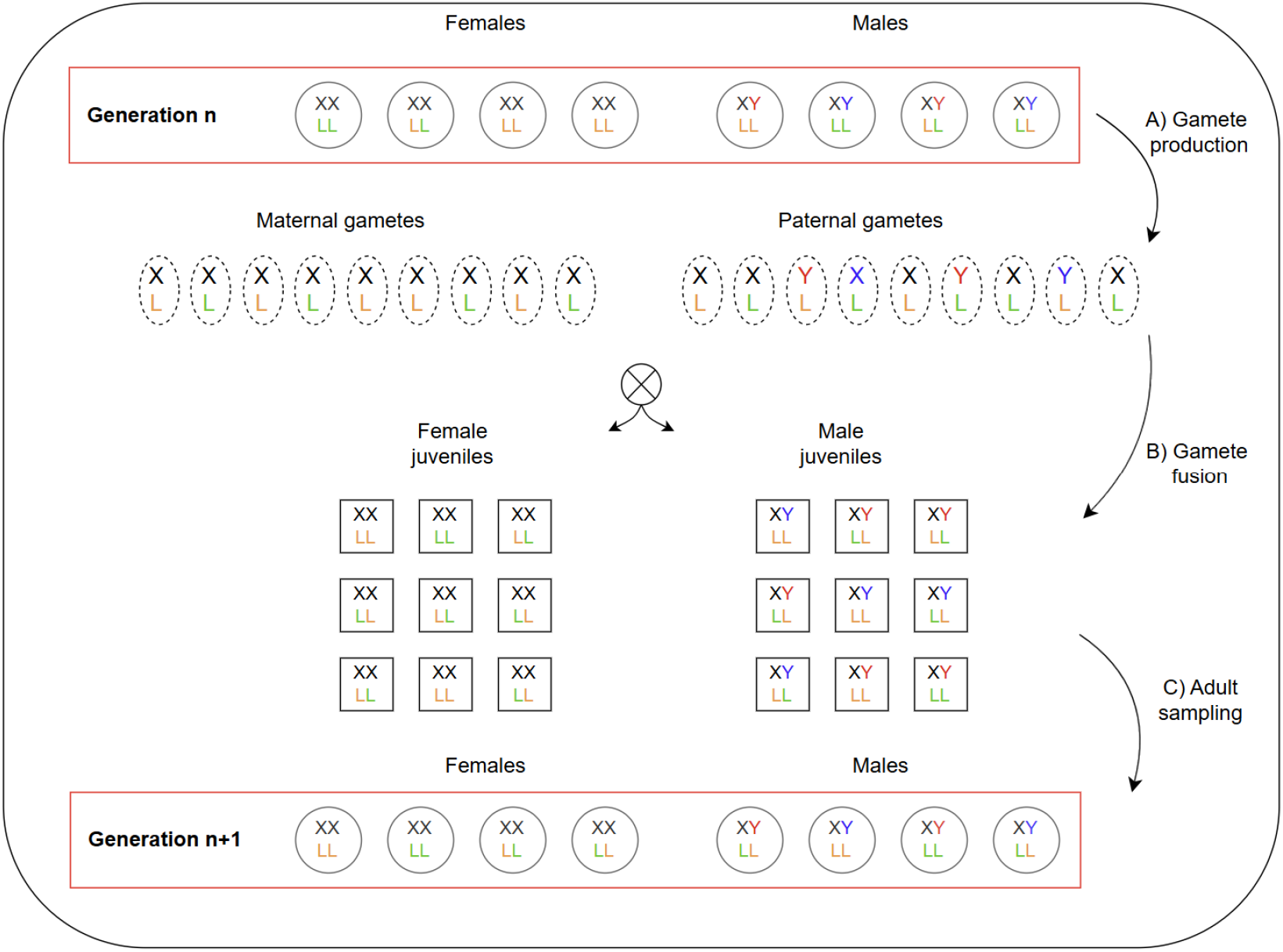
Illustration of the reproduction step between one generation and the next in our two-sex diploid Wright-Fisher model with selection. For each gamete or individual, we represent the X and Y chromosomes and an autosomal locus L. The autosomal locus can carry two alleles (orange and green) and so does the Y chromosome (red and blue). An infinite number of juveniles from each sex are created by fusing gametes of different genotypes. These gametes are created in proportions that depend on the allele frequency in the parental generation (see main text). Juveniles are then separated between those that received a paternal Y chromosome and those that received a paternal X chromosome. Finally, *N/*2 individuals are drawn from each subgroup to form the next adult generation, ensuring that the sex ratio remains balanced. This schematic representation encompasses the two chromosomal contexts of interest: if the focal mutation is located on the Y chromosome, then the alleles at the autosomal locus have no impact on the juvenile sampling probabilities (as only the mutant or ancestral allele carried by Y chromosome affects the selection coefficient); if the mutation is carried by an autosome, then the Y-chromosome alleles do not affect the juvenile sampling probabilities (as selection is supposed to only act on locus L).

### 2.2 One-step change in frequencies

We start by studying how the frequency of the mutation changes from one generation to the next. To compute these one-step transitions, we take the specificities of the two contexts (*i*.*e*., mutations on autosomes and on Y chromosomes) into account.

#### Mutation on an autosome

Let us first argue that, in this context, the mutation frequency in the male and female subpopulations can be considered equal in each generation. Denoting *p*^*m*^ (*resp., p*^*f*^ ) the mutation frequency among males (*resp*., females), the gametes with associations X*A*, X*a*, Y*A* and Y*a* produced by males have frequencies given by (1 − *p*^*m*^)*/*2, *p*^*m*^*/*2, (1 − *p*^*m*^)*/*2 and *p*^*m*^*/*2. Indeed, in this context we assume no linkage between the mutation and the X and Y chromosomes. Each female transmits an X chromosome associated with the mutation with frequency *p*^*f*^ . Merging all these gametes, we obtain male and female juveniles carrying each genotype *AA, Aa* and *aa* with identical frequencies, given by

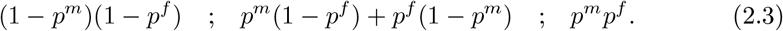

The *N/*2 male and female adults are then drawn from this juvenile population. Since selection acts similarly in the two sexes, this mechanism ensures that the frequencies of the mutation among male and female adults in the new generation are equally distributed. This is true even if *p*^*m*^ and *p*^*f*^ are different. Because we consider population sizes of at least 100 individuals, the number of males and females carrying each genotype is close to its expectation, and we can therefore make the approximation that the mutation frequencies are equal in the two sexes in each generation. Writing *p* = *p*^*m*^ = *p*^*f*^ for this shared frequency, the juvenile frequencies in (2.3) can be rewritten (1 − *p*^2^), 2*p*(1 − *p*) and *p*^2^.

Next, let *p*_*n*_ ∈ [0, 1] denote the frequency of the mutant allele in the autosomal population in the *n*th generation. That is,

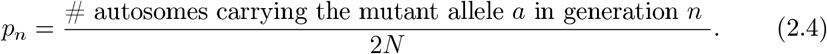

The genotype frequencies in the juvenile population weighted by the corresponding selection coefficients are given by :

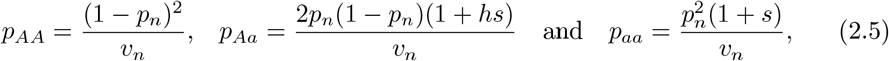

with

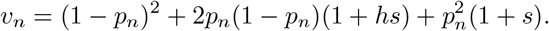

This matches the expression (2.2) with males and females grouped. Conditionally on *p*_*n*_, the frequency of the mutant allele *a* in the next generation is therefore given by

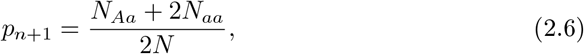

where

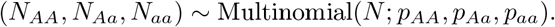

The multinomial law corresponds to the fact that each of the *N* adults inherits a genotype *AA, Aa* and *aa* independently with probabilities *p*_*AA*_, *p*_*Aa*_ and *p*_*aa*_ defined in (2.5).

#### Mutation on a Y chromosome

We now consider that the mutation *a* appears on the Y chromosome at a locus that does not recombine with the sex-determining locus, so that this mutation is never transferred onto an X chromosome. It is therefore enough to keep track of the mutation on the Y chromosomes only (*i*.*e*., among the male sub-population). The form of the selection coefficients remains identical to the autosomal case for the two feasible genotypes *AA* and *Aa*, and we never observe the genotype *aa*.

Let *q*_*n*_ denote the frequency of the mutation *among the Y chromosome sub-population*. That is,

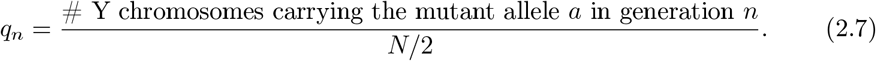

Note that the denominator is now *N/*2, which corresponds to the total number of Y chromosomes in the population. Each adult male is of genotype *AA* or *Aa* with respective probabilities

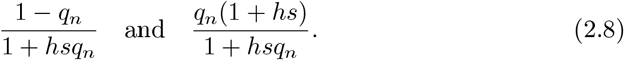

This matches the haploid Wright-Fisher model with selection described for instance in Chapter 5.2 in (Etheridge, 2011). The frequency of the mutation among the *N/*2 Y chromosomes in the next generation is thus given by *q*_*n*+1_ = 2*N*_*a*_*/N*, where

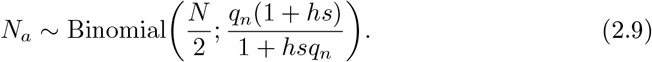

Focusing on the Y chromosome sub-population by following the dynamics of *q*_*n*_ defined in (2.7) instead of following the mutation frequency at the population level (as in the definition of *p*_*n*_ in (2.4)) means that fixation can be studied in the same way as in the autosomal case. Indeed, if we were to consider the whole chromosomal population, the mutation frequency would be capped at 0.25. Here, both (*p*_*n*_)_*n*∈ℕ_ and (*q*_*n*_)_*n*∈ℕ_ take their values in [0, 1], where 0 and 1 are absorbing states corresponding respectively to extinction and fixation. The appropriate choice for the initial frequencies *q*_0_ relatively to *p*_0_ to account for the difference in effective population size between autosomes and Y chromosomes is provided in Section 2.4.

### 2.3 Diffusion approximations

We wish to investigate the dynamics of the frequency of the mutation in the population through time. In our model, this frequency is modelled by the discrete-time Markov chains (*p*_*n*_)_*n*≥0_ and (*q*_*n*_)_*n*≥0_, taking values in

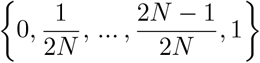

for autosomal mutations and in

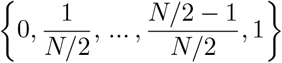

for mutations on the Y chromosome. Although we can easily compute their Markov transition kernel, a direct analysis of their long-term dynamics is practically impossible for biologically relevant values of *N*, as the transition matrices contain of the order of *N* ^2^ non-zero coefficients and computing interesting quantities such as fixation probabilities would require to solve a prohibitive number of equations. Nevertheless, as in the classical Wright-Fisher framework we can perform a large-population approximation of these discrete-time Markov chains using diffusion processes, as shown below.

For any *N* ∈ ℕ, we write *s*^*N*^ = *s*_0_*/N* with *s*_0_ ∈ ℝ and we define the processes 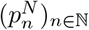 and 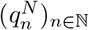 as in Section 2.2, with parameters values *h* and *s*^*N*^ . They represent the random trajectories of the frequency of a mutation appearing on an autosome and on a Y chromosome, respectively, in a population of *N* diploid individuals. Let us write ⌊*x*⌋ for the integer part of a real number *x* and let *B* denote the standard Brownian motion. Rescaling time by a factor *N*, that is, considering a new timescale where one unit of time equals *N* generations, the rescaled processes 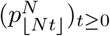 and 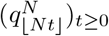 respectively converge as *N* → ∞ to the solutions to the equations:

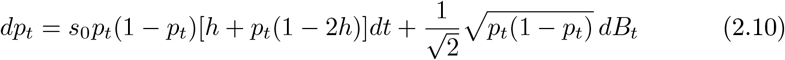

for autosomal mutations, and

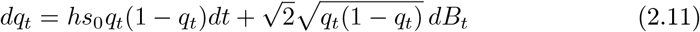

for mutations on the Y chromosome.

From now on, we write (*p*_*t*_)_*t*≥0_ and (*q*_*t*_)_*t*≥0_ for the continuous solutions to these equations, that is, the diffusions respectively representing the frequency of the mutation which appeared on an autosome and on a Y chromosome. The exact nature of this convergence, as well as a rigorous proof, can be found in Appendix A.3.

Diffusions are classically composed of two parts: a component giving the general tendency to increase or decrease, called the *diffusion drift*, and a term encoding the mean-zero fluctuations around the general tendency, called the *variance term*. This nomenclature typically causes some confusion, as the drift term of the diffusion corresponds to the effect of selective forces, and the variance due to the randomness in reproduction corresponds to the effect of *genetic drift*. Therefore, we will always specify *genetic drift* when referring to this source of randomness, and use the term *diffusion drift* when talking about the mathematical drift term.

We can first observe that the diffusion (*q*_*t*_)_*t*≥0_ introduced in (2.11) exhibits a variance coefficient twice as large as that of (*p*_*t*_)_*t*≥0_ (2.10). This difference arises from the increased genetic drift affecting Y chromosomes, whose effective population size corresponds to a fourth of the autosomal effective population size. In the Y chromosome context, only heterozygotes can have a disadvantage or an advantage and thus, for fixed *h* and *s*_0_, the diffusion drift of (*q*_*t*_)_*t*≥0_ always remains of the same sign. By contrast, the diffusion drift of (*p*_*t*_)_*t*≥0_ is more complex and can change sign depending on the mutation frequency.

One-dimensional diffusions are particularly suitable for our study, as many theoretical results have been developped to analyse their stochastic behaviour (see, *e*.*g*., Chapter 7 in Durrett 2008 or Chapter 3 in Etheridge 2011). In the cases we consider, it is possible to find explicit expressions for the probability of absorption in 0 or 1, corresponding to the purging and the fixation of the mutation, respectively. We also know that one of these two events will happen in finite time with probability 1 (Ikeda and Watanabe, 1989), and we can compute the average time for the mutation to be fixed or removed from the population, *i*.*e., the mutation mean segregation time*. It is important to note that we obtain our limiting diffusions on precisely the same timescale (*Nt, t* ≥ 0) in both chromosomal contexts (autosomes and Y chromosomes), which will allow us to compare the mean segregation times on autosomes and on Y chromosomes in a straightforward way.

Diffusions of the form taken by (*p*_*t*_)_*t*≥0_ and (*q*_*t*_)_*t*≥0_ have been known and studied for a long time. Diffusion approximations for the trajectories of allele frequencies trace back to Kimura (1962), who first computed the fixation probability of a favourable allele in a haploid Wright-Fisher model with selection using a diffusion similar to (*q*_*t*_)_*t*≥0_. The process (*p*_*t*_)_*t*≥0_ emerges from asexual diploid Wright-Fisher models (Durrett 2008, chap. 7; Etheridge 2011, chap. 5), and was also used in Takahata (1990) in the special case of balancing selection (*h* < 0, *s*_0_ < 0).

In our framework, both (*p*_*t*_)_*t*≥0_ and (*q*_*t*_)_*t*≥0_ are derived from the same population dynamics, the same selective coefficients and on the same time scale. The differences between them come from the *structural effects* of the heterogametic Y chromosome as opposed to the autosomes, namely the set of feasible genotypes and the smaller population size inducing stronger genetic drift. We will therefore be able to study how these differences affect the mutation fixation probability and mean segregation time by directly comparing (*p*_*t*_)_*t*≥0_ and (*q*_*t*_)_*t*≥0_.

### 2.4 Deriving mean segregation times and fixation probabilities

As described in the previous section, we can derive exact formulae for the fixation probabilities and mean times to purge or fixation, given the dominance parameter *h*, the selection coefficient *s*_0_ and initial mutation frequencies *p*_0_ and *q*_0_.

For *i* = 0 (*i*.*e*., extinction) or *i* = 1 (*i*.*e*., fixation), we write 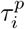 and 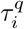 for the random times

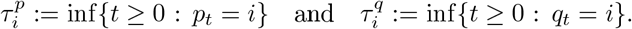

They correspond to the times at which each diffusion reaches one of the absorbing states 0 or 1. Denote by 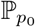 (*resp*., 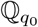) the probability measure governing (*p*_*t*_)_*t*≥0_ started at *p*_0_ ∈ (0, 1) (*resp*., (*q*_*t*_)_*t*≥0_ started at *q*_0_ ∈ (0, 1)). We first compute the fixation probability of the mutation in each context. All the details can be found in Appendix A.3.

In the autosomal context, we obtain:

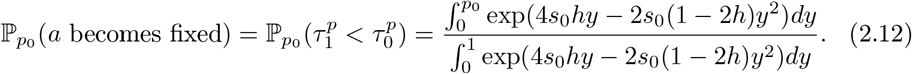

This is an explicit value, although it is difficult to write it in a simpler way to analyse its behaviour as a function of *h* and *s*_0_. The value for the mutation fixation probability in the Y chromosome context is simpler and well known (Crow and Kimura, 1970):

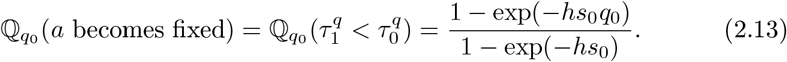

Since the diffusion necessarily hits one of the boundaries 0 or 1 in finite time, fixation and extinction are complementary events and

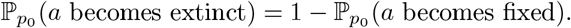

The expressions for the mean time to purge or fixation are more complex and are detailed in Appendix A.3. It is important to observe that these quantities are the mean times to reach *any* of the boundaries, be it 0 or 1. Mathematically speaking, they are the mean absorption times of the processes (*p*_*t*_)_*t*≥0_ and (*q*_*t*_)_*t*≥0_. Biologically speaking, they correspond to the average segregation time of the mutation in the population (before it becomes fixed or extinct).

Although the population size *N* does not appear in the limiting diffusions, and therefore in the fixation probability and mean segregation time formulae, we can still adjust the initial frequencies to approximate the effects of various population sizes. For example, considering that only a single mutant copy can initially be found in a population of *N* individuals, then the appropriate initial frequency is 2*/N* in the Y chromosome context and 1*/*(2*N* ) in the autosome context.

### 2.5 Individual-based simulations

The method based on diffusion approximations that we propose enables a fair comparison of the fate of mutations on Y chromosomes versus autosomes in our model, carefully considering the same timescale in both contexts. However, it remains necessary to test the appropriateness of this timescale and to evaluate the precision of our analytical large-population approximation. To do so, we performed individual-based simulations using SLiM 4.3. We simulated a single panmictic population of size *N* = 100, *N* = 1000 or *N* = 10, 000 diploid individuals under a Wright–Fisher model. Each individual carried two pairs of chromosomes, both assumed to be non-recombining along their entire length. The first pair corresponded to the sex chromosomes and included a sex-determining locus with two alleles, one of which was permanently heterozygous, mimicking a classical XY sex-determination system. Individuals therefore carried either XY chromosomes (males) or XX chromosomes (females). The second pair corresponded to autosomes. For each parameter set, we performed 10,000 replicate simulations in which a single copy of a mutation was introduced into a randomly chosen individual, either on the autosome or on the Y chromosome. Mutations arising on the Y chromosome were therefore completely linked to the male-determining allele (recombination rate = 0), whereas mutations arising on autosomes recombined freely with the sex-determining locus (recombination rate = 0.5). Simulations were run until the mutation was either fixed or lost, with a maximum duration of 100,000 generations.

We compared the results of the simulations to the theoretical results obtained with the approximations (*p*_*t*_)_*t*≥0_ and (*q*_*t*_)_*t*≥0_ using the same sets of parameters. As mentioned earlier, the population *N* does not appear directly in the diffusions but still plays two important roles in our comparisons. First, we use the initial frequencies *p*_0_ = 1*/*(2*N* ) and *q*_0_ = 2*/N*, as explained in Section 2.4. Second, as the diffusion approximations were obtained as limits of processes where one unit of time corresponded to *N* generations, we can recover the original time units by multiplying the diffusion timescale by *N* to compare the analytical predictions with the simulations.

## 3 Results

To better understand the fate of mutations on the Y chromosome and on autosomes, we proceed in two steps. First, we analyse the diffusions obtained in the Y chromosome and autosome contexts separately, focusing on how the fixation probability and mean segregation time of the mutation vary with the parameters *s*_0_ and *h*. We analyse the dynamics of alternative types of mutations: strictly beneficial, strictly deleterious, underdominant and overdominant (see Table 1). Second, we compare these statistics between the Y chromosome and autosome contexts to better understand how the difference in population size and heterozygosity affects the dynamics of the mutation allele frequency.

**TABLE 1:**
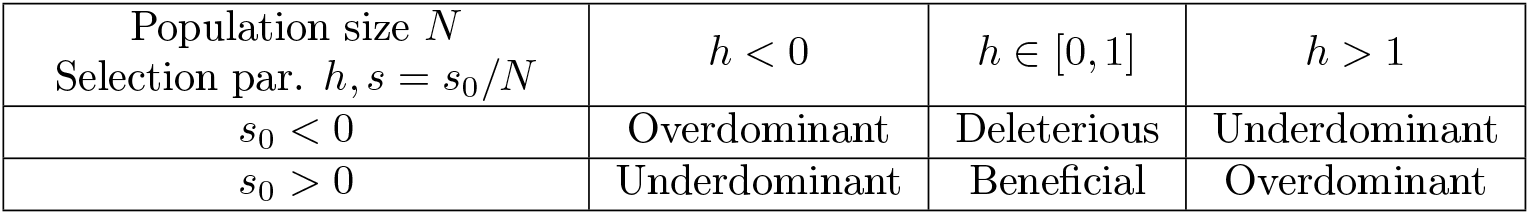
Description of the different selective scenarios in the autosomal context, according to the values of the selection parameters *h* and *s*_0_ which induce differences in fitness given by 1 for the genotype *AA*, 1+*hs*_0_*/N* for the genotype *Aa* and 1+*s*_0_*/N* for the genotype *aa* (where *N* stands for the population size). In the Y chromosome context, the mutation never appears at homozygous state and it can thus be only beneficial (*hs*_0_ > 0) or deleterious (*hs*_0_ < 0).

| Population size $N$<br>Selection par. $h, s = s_0/N$ | $h < 0$ | $h \in [0, 1]$ | $h > 1$ |
| --- | --- | --- | --- |
| $s_0 < 0$ | Overdominant | Deleterious | Underdominant |
| $s_0 > 0$ | Underdominant | Beneficial | Overdominant |

### 3.1 Mutation on a Y chromosome

The interpretation of the results for the Y chromosome context is straightforward. The fixation probability and mean segregation time (*i*.*e*., the mean absorption time in 0 or 1 of (*q*_*t*_)_*t*≥0_) only depend on the product *hs*_0_. Figure 2 shows the fixation probability and mean absorption time of the process for different values of *hs*_0_, ranging from scenarios where the mutation is deleterious (*hs*_0_ < 0) to scenarios where it is beneficial (*hs*_0_ > 0). As expected, the probability of fixation increases when *hs*_0_ increases. When *hs*_0_ is close to 0, the probability of fixation of the mutation is given by its initial frequency, which is expected from the neutral haploid Wright-Fisher model.

**FIGURE 2:**
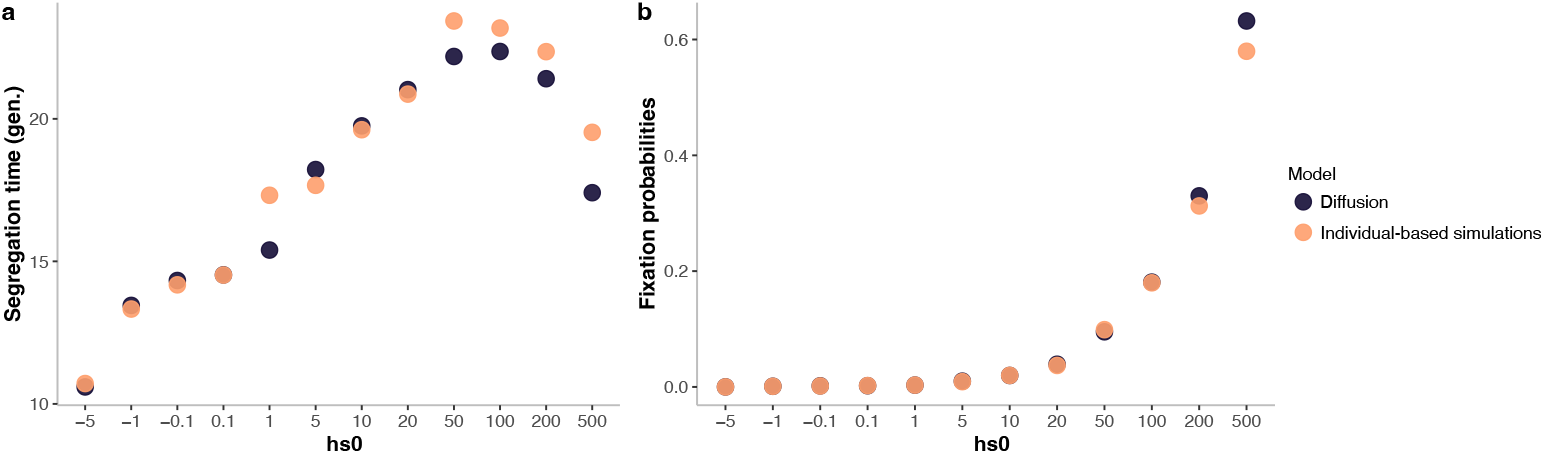
Mean segregation times (**a**) and fixation probabilities (**b**) obtained from the diffusion (*q*_*t*_) (2.11) started at *q*_0_ = 2*/N*, with *N* = 1000. For each pair of parameters *h* and *s*_0_, they approximate the mean segregation time and fixation probability of a single mutation appearing on a Y chromosome of an individual within a population of size *N*, where the selection parameters induce differences in fitness given by 1 for the genotype *AA* and 1 + *hs*_0_*/N* for the genotype *Aa* in accordance with (2.1). The mean segregation time is expressed in terms of number of generations (denoted *gen*. in the Figure). For each value of the product *hs*_0_, the black dots correspond to the results obtained with our diffusion approximations, and the orange dots to the results of the individual-based simulations described in Section 2.5.

The mean segregation time is non-monotonic. When *hs*_0_ is negative, the fixation probability is close to 0 and the mutation quickly disappears from the population. Increasing *hs*_0_ above 0 increases the probability that the mutation goes to fixation, which takes on average longer than the purge of a mutation and thus increases the mean segregation time. When *hs*_0_ becomes large (*hs*_0_ > 100 for *N* = 1000), mutation fixation further accelerates, thereby reducing the mean segregation time. It therefore appears that beneficial mutations with intermediate positive selective coefficients are those with the largest average segregation time on Y chromosomes. Individual-based simulations confirmed the accuracy of our diffusion approximation, even for populations as small as *N* = 100 (see also Appendix A.2). They also show that our approach via diffusion approximations allows us to estimate very small but nonzero fixation probabilities across certain regions of the parameter space, where individual-based simulations fail to provide reliable estimates due to the rarity of fixation events.

### 3.2 Mutation on an autosome

The behaviour of mutations on autosomes is more complex, as homozygotes and heterozygotes undergo different and potentially opposite selective pressures. We therefore examine the four possible types of mutations described in Table 1 separately: beneficial, deleterious, overdominant and underdominant.

#### Beneficial mutation

When 0 < *h* < 1 and *s*_0_ > 0, the mutation is strictly beneficial and the diffusion drift term of (*p*_*t*_)_*t*≥0_ remains positive at any time, mainly because the selective advantage of the mutation is primarily conferred by homozygotes. The resulting fixation probabilities and segregation times are shown in Figure 3. In brief, as in the Y chromosome context, increasing *s*_0_ leads to higher fixation probabilities, and the mean segregation time is non-monotonic.

**FIGURE 3:**
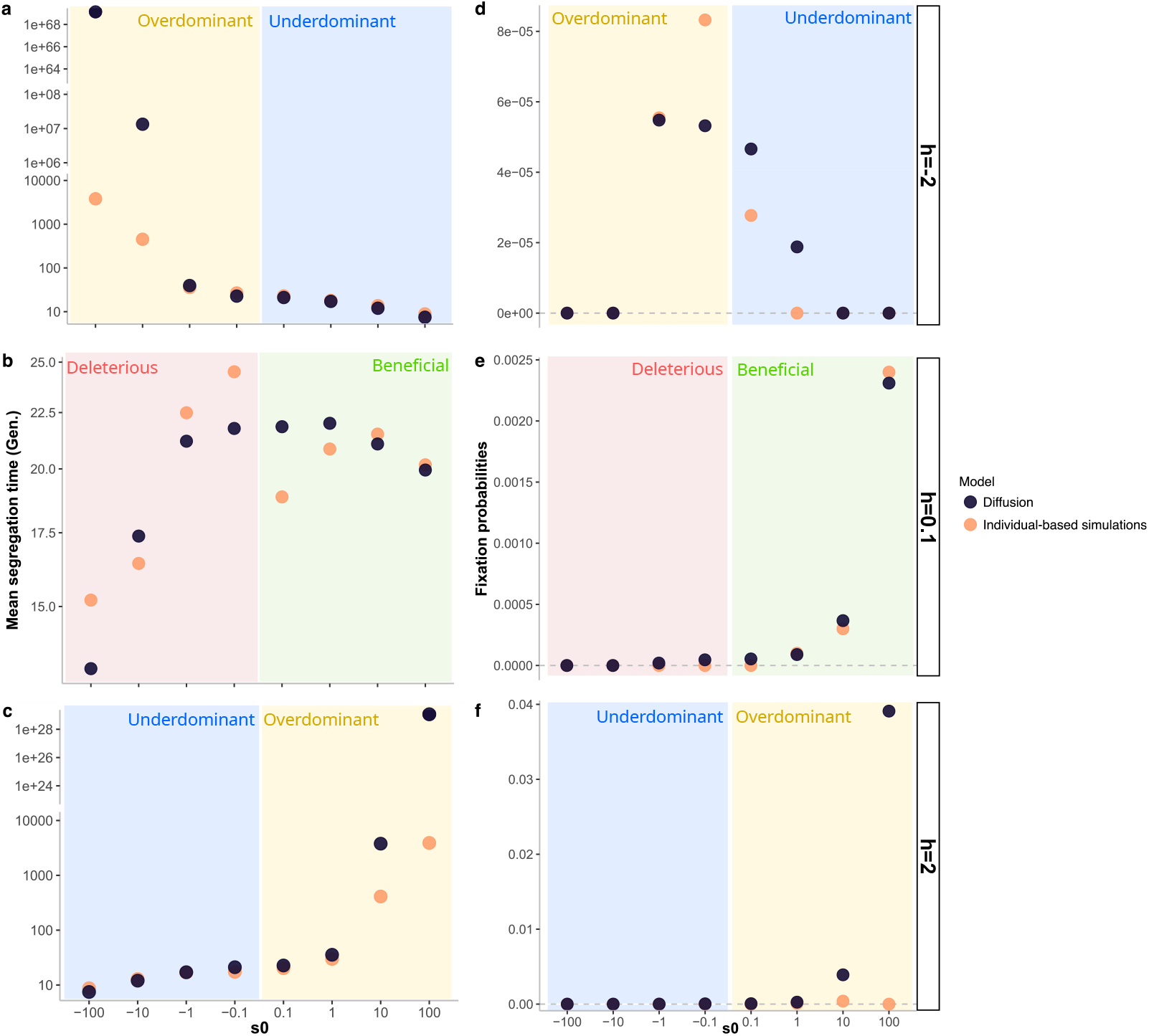
Mean segregation times (**a, b** and **c**) and fixation probabilities (**d, e** and **f** ) obtained from the diffusion (*p*_*t*_)_*t*≥0_ (2.10) started at *p*_0_ = 1*/*(2*N* ), with *N* = 10000. For each pair of parameters *h* and *s*_0_, they approximate the mean segregation time and fixation probability of a single mutation appearing on an autosome of an individual within a population of size *N*, where the selection parameters induce differences in fitness given by 1 for the genotype *AA*, 1+*hs*_0_*/N* for the genotype *Aa* and 1+*s*_0_*/N* for the genotype *aa* in accordance with (2.1). The mean segregation time is expressed in terms of number of generations (denoted *gen*. in the Figure). Each panel presents the results obtained for different values of *s*_0_ and fixed *h* (*h* = − 2 in panels **a** and **d**, *h* = 0.1 in panels **b** and **e**, *h* = 2 in panels **c** and **f** ); the black dots correspond to the results obtained with our diffusion approximations, and the orange dots to the results of the individual-based simulations described in Section 2.5. The four scenarios described in Table 1 are made explicit by the coloured backgrounds (green for beneficial, red for deleterious, yellow for overdominant and blue for underdominant). The mean segregation times in the individual-based model and in the diffusion approximation mainly differ when the diffusion approximation predicts segregation times that can potentially be much larger than the 100,000 generation cut-off used in the simulations; likewise, the differences in fixation probabilities appear in situations where the fixation probability is too small to be correctly inferred from 10,000 simulations.

Increasing *h* also increases the fixation probability, because heterozygotes—which dominate when the mutation is rare—benefit from a stronger positive selective advantage, helping the mutation to avoid extinction early on. Higher fixation probabilities also result in longer mean segregation times, as mutations that eventually fix tend to persist longer at intermediate frequencies than those which eventually disappear. Figure 3 shows a close agreement between our results and the simulations for beneficial mutations.

Two cases are of analytical and biological interest: *h* = 0 and *h* = 1*/*2. When *h* = 1*/*2, the mutation is co-dominant and its fitness advantage is equally conferred by homozygotes and heterozygotes. In that case, the large-population diffusion approximation is given by:

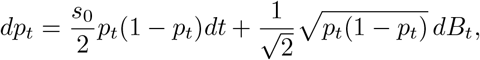

and the associated diffusion drift term exactly matches that in the equation describing the mutant allele frequency in the Y chromosome context (2.11). We can therefore directly compute the fixation probability on autosomes which is given by

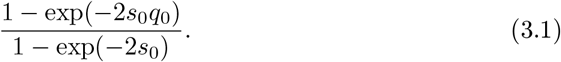

When *h* = 0, the mutation is completely recessive and only the homozygotes *aa* benefit from a selective advantage. The diffusion approximation becomes:

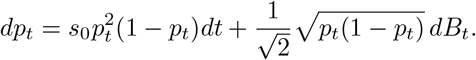

In this equation, the diffusion drift resembles the diffusion drift term computed in the Y chromosome context (2.11), but multiplied by a factor *p*_*t*_. Indeed, when *p*_*t*_ is close to 0 and the mutation not widespread in the population, the homozygotes *aa* which actually benefit from a selective advantage are rare, and the factor *p*_*t*_ substantially reduces the diffusion drift term. On the contrary, when *p*_*t*_ →1, homozygotes *aa* predominate and the added factor *p*_*t*_ has no impact.

#### Deleterious mutations

When 0 < *h* < 1 and *s*_0_ < 0, the mutation is deleterious and its diffusion drift is always negative. Fixation probabilities are therefore small (strictly less than 1*/*2*N* ) and mean mutation segregation times are short (a few generations). These quantities decrease when increasing |*s*_0_|, as expected (see panels **b** and **e** in Figure 3). As before, individual-based simulations confirmed the accuracy of our diffusion approximation.

#### Overdominant mutations

When *h* < 0 and *s*_0_ < 0 or when *h* > 1 and *s*_0_ > 0, the mutation behaves as over-dominant, meaning that the heterozygotes *Aa* have a fitness advantage over the two homozygotes *AA* and *aa*. This leads to negative frequency-dependent balancing selection, in which selection tends to maintain the mutation at an intermediate frequency. More precisely, such cases lead to the existence of *metastable states*, which are particular frequencies in (0, 1) that cancel the diffusion drift term, and close to which the stochastic fluctuations due to genetic drift are the main drivers of allele frequency changes. It is straightforward to compute the value of the metastable state:

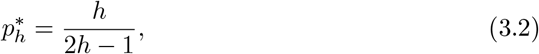

in line with classical results (Lefevre et al., 2016). This frequency 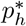 is *attractive* as selection always pushes the mutation frequency towards it: the diffusion drift is positive when 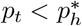 and negative when 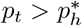 . Eventually, the random fluctuations due to genetic drift cause the mutant allele frequency to deviate far enough from the metastable state for the frequency to reach 0 or 1, *i*.*e*., for the mutation to become extinct or fixed. This can lead to extremely large segregation times, especially when *N* is large, as the frequency will typically oscillate around 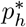 for a very long time until a rare stochastic excursion drives it away (see panels **a** and **c** in Figure 3). The mean segregation time depends on both the strength of selection conferred by *s*_0_ and the distance of the metastable frequency from 0 or 1, which is determined solely by *h*. In general, mean segregation times increase as |*h*| deviates further from zero. For large values of *h*, the metastable state approaches 0.5 and the fixation probability can become close to 0.5. Indeed, once the mutation frequency has reached the metastable state, the mutation has nearly equal probabilities of ultimately fixing or being lost through long stochastic fluctuations around the metastable frequency. However, in a large population, fixation events may occur only on a virtually unobservable timescale. For instance, with *N* = 1000, *h* = 5 and *s*_0_ = 100, mutations will segregate during 1.0*e*193 generations on average.

In such cases, the diffusion approximation yields values that may substantially differ from those obtained in individual-based simulations. This discrepancy arises because we stopped our simulations after 100,000 generations, even if fixation or loss has not occurred yet, which underestimates both the fixation probabilities and the mean segregation time. For example, when *N* = 10, 000, *h* = −10 and *s*_0_ = −1, the theoretical fixation probability of the mutation is 0.00025. Out of 10,000 simulations, we would therefore expect to observe approximately 2.5 fixation events. Yet, while fixation never occured in the simulations, in 15 of them the mutation was still segregating after 100,000 generations, suggesting that some of them may eventually fix and we are simply not able to observe it within the time window considered. This phenomenon highlights the value of our theoretical approach, as reaching definitive conclusions from simulations alone is difficult in these cases.

These considerations illustrate one of the key messages of this study, namely that the fixation probability alone, without consideration of the mean time to fix or go extinct, is not enough in general to predict the fate of a mutation.

#### Underdominant mutations

When *h* > 1 and *s*_0_ < 0, the mutation is underdominant, meaning that the heterozygotes have a lower fitness than both homozygotes. In this case, even if the mutation has a positive selective advantage at homozygote state, it has a low fixation probability, since the counter-selection it experiences at the heterozygous state hinders its spread when rare and leads to its rapid purge (see panels **c** and **f** in Figure 3). By contrast, such mutations rapidly fix when they become abundant, as the decreasing relative frequency of heterozygotes increases the relative fitness of the mutation. Similarly to the overdominant scenario, the diffusion drift is canceled at the frequency *h/*(2*h* −1). However, in this case the diffusion drift is negative below the critical frequency and positive above it. This frequency therefore does not constitute a metastable state; it is *repulsive* as selection tends to drive the mutation frequency away from this value. Underdominant mutations thus have a short mean segregation time (*e*.*g*., 12 generations for *h* = 2 and *s*_0_ = −10), similar to deleterious mutations over most of the parameter space.

When *s* < 0 and *h* > 1, the mutation is also underdominant, but this time it is deleterious for both heterozygotes and homozygotes *aa*. Accordingly, its frequency dynamics closely resemble those of deleterious mutations.

### 3.3 Comparing the mutation dynamics on autosomes and on Y chromosomes

We now turn to a direct comparison of the mutation fixation probabilities and mean segregation times on the Y chromosome and on autosomes. We perform these comparisons using identical sets of parameters, covering all the cases exposed in Table 1. This allows us to isolate the effects of the specific transmission of alleles through the Y chromosome, comparatively to autosomes. In particular, since both processes (*p*_*t*_)_*t*≥0_ and (*q*_*t*_)_*t*≥0_ evolve on the same time scale, we can directly contrast the mean times of interest.

To enable meaningful comparisons between the autosome and Y chromosome context, it is essential to account for the fact that autosomes experience an effective population size four times larger than Y chromosomes. Since our focus is on the fate of a mutation *after it has arisen* (rather than on its appearance rate), we use initial allele frequencies that scale accordingly: *p*_0_ = 1*/*(2*N* ) for the autosome and *q*_0_ = 2*/N* for the Y chromosome. These differences in initial frequencies influence both the fixation probabilities and the mean segregation times in ways that depend on the values of *s*_0_ and *h*. Importantly, this framework does not capture the fact that, in a diploid population with balanced sex ratio, mutations arise more frequently on autosomes than on the Y chromosome; our results therefore reflect fixation dynamics rather than substitution rates. Nevertheless, the approach is flexible and could readily accommodate alternative assumptions about the initial allele frequency, including equal starting frequencies across genomic locations or scenarios where the mutation has already reached a substantial frequency.

Figure 4 shows the fixation probability and the mean segregation time for mutations appearing on a Y chromosome or an autosome, as a function of the selection parameters *h* and *s*_0_. In particular, Figure 4**b** displays the ratio of fixation probabilities between the Y chromosome and autosomes in order to illustrate in which case fixation is more likely. Charlesworth et al. (1987) studied a similar ratio but considering substitution rates, which adds a factor 4 to the ratio. This factor arises from the fact that, assuming that the mutation rate is the same for each individual chromosome, mutations are four times as frequent on autosomes as on Y chromosomes since their population size is four times larger. We can study a similar classical substitution rate comparing the results of Figure 4**b** to the threshold value of 4. For example, in the purely neutral case, the fixation probability is 4 times higher on Y chromosomes, resulting in equal substitution rates between the two contexts. However, our parametrization differs from that of Charlesworth et al. (1987) and Figure 4**b** cannot be used directly to assess concordance with our results. A precise comparison is provided in Appendix A.1. Charlesworth et al. (1987) studied the substitution rates in the Y chromosome and on autosomes with either beneficial or deleterious mutations. For beneficial mutations, they used Haldane’s branching approximation and our results match theirs whenever *s*_0_ is sufficiently large (*e*.*g*., whenever *hs*_0_ > 10 for *N* = 10, 000). However, Haldane’s approximation breaks down as *s*_0_ decreases toward near-neutrality (*e*.*g*., whenever *s*_0_ < 10 for *h* = 0.1), whereas our method still yields expected values close to 0.25 (see Table A.1 in Appendix A.1). In the deleterious scenario, Charlesworth et al. (1987) used the same diffusion approximation as we do but performed additional Taylor expansions to derive simpler analytical values for the quantities appearing in (2.12) and (2.13). Our results coincide with theirs when these expansions are possible. Our approach therefore provides a unified tool to study a continuum of scenarios across a wide range of parameters. Figure 4**b** displays the mean segregation time of the mutation in the two contexts. As discussed in the overdominant scenario, this time information is crucial here as it strongly affects the number of fixed mutations observed within a given time window. We now turn to an analysis across the entire parameter space.

**FIGURE 4:**
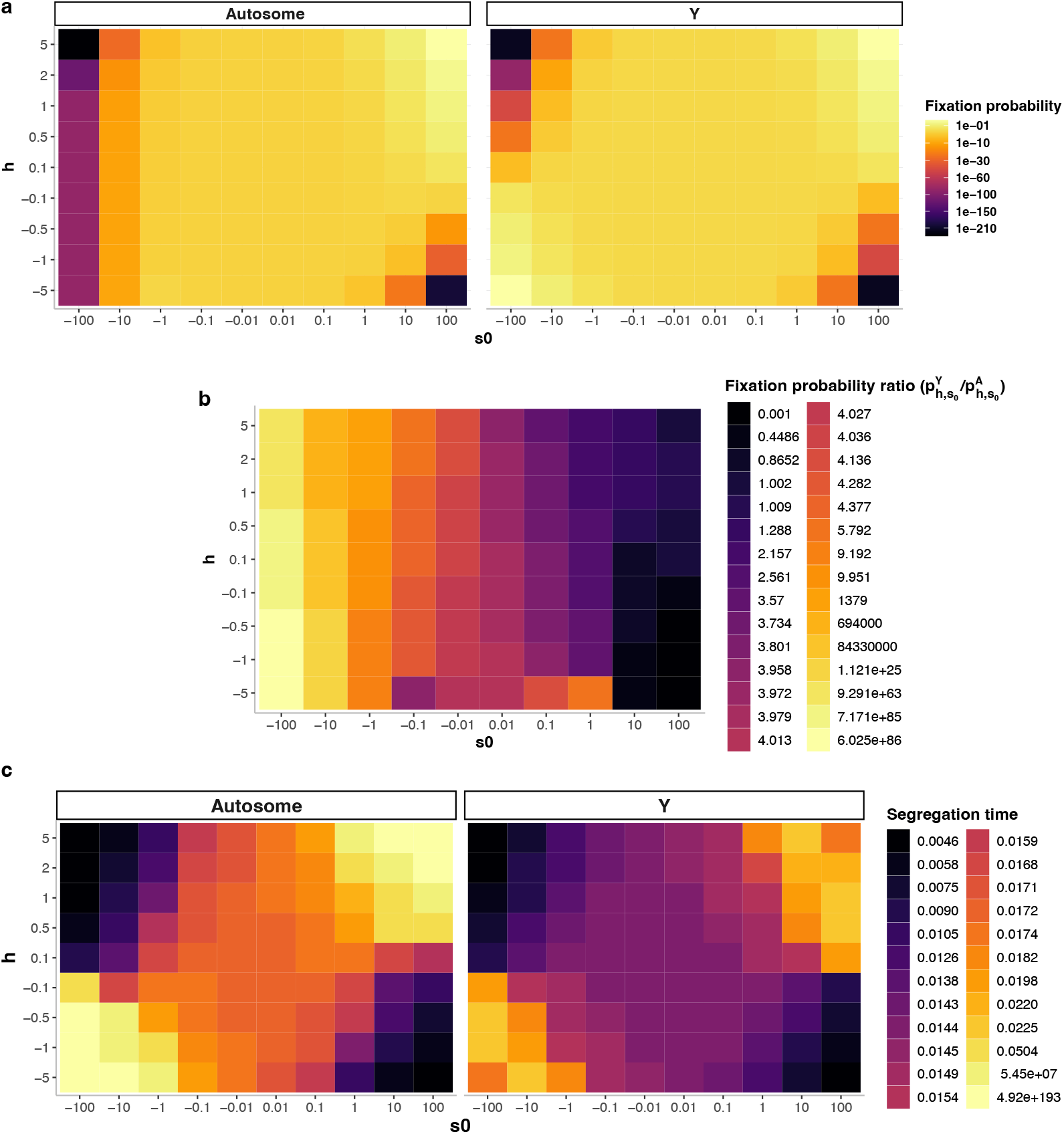
Fixation probabilities (**a**) and mean segregation times (**c**) obtained from the diffusion (*p*_*t*_) (2.10) started at *p*_0_ = 1*/*(2*N* ) (left) and from the diffusion (*q*_*t*_) (2.11) started at *q*_0_ = 2*/N* (right), with *N* = 1000. For each pair of parameters *h* and *s*_0_, they approximate the mean segregation time and fixation probability of a single mutation appearing on an autosome of an individual within a population of size *N*, where the selection parameters induce differences in fitness given by 1 for the genotype *AA*, 1 + *hs*_0_*/N* for the genotype *Aa* and 1 + *s*_0_*/N* for the genotype *aa* in accordance with (2.1). The mean segregation times are expressed in the diffusion timescale and can be multiplied by *N* to recover a scale in generations. Panel **b** displays, for each parameter set (*h, s*_0_), the ratio of the fixation probability on Y chromosomes 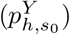 and that on autosomes 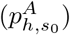. When this ratio is above 1, a single mutation is more likely to fix on the Y chromosome than on autosomes. When it is above 4, the *substitution rate* (see main text) is higher on the Y chromosome than on autosomes.

First, advantageous mutations (*h* > 0, *s*_0_ > 0) can have a higher probability of fixation on autosomes than on Y chromosomes but only when they are beneficial and not overdominant, according to the classification in Table 1 (see, *e*.*g*., the case *s*_0_ = 10, *h* = 0.1 in Figure 4**b** This is expected, as beneficial mutations with a moderate selective advantage are more likely to fix when genetic drift is weaker, as the latter counteracts selection. This effect is particularly pronounced when *h* is close to 0, as recessive mutations have a reduced selective advantage (and therefore a lower fixation probability) on the permanently heterozygous Y chromosome. Consistent with the nearly neutral theory (Ohta, 1973), slightly beneficial mutations (0 < *s*_0_ ≤ 1, 0 < *h* < 1) exhibit a higher fixation probability on the Y chromosome, as their nearly neutral behavior makes them more likely to fix in smaller populations (see Figure 4). Fixation is also more likely on Y chomosomes when the strength of selection is strong (*s*_0_ ≥ 100), as genetic drift becomes negligible and the mutation is again more likely to fix in the smaller Y chromosome population.

When the mutation is underdominant and advantageous at homozygote state (*h* < 0 and *s*_0_ > 0, see Table 1), fixation probabilities on autosomes can also exceed those on the Y chromosome (*e*.*g*. when *h* = − 1 and *s*_0_ = 100). This occurs because these mutations are always deleterious on the Y chromosome, unlike on autosomes where homozygous individuals *aa* benefit from a selective advantage. The smaller effective population size of the Y chromosome may also compensate for this difference at smaller parameter values (*e*.*g*., in the case *h* = − 0.1 and *s*_0_ = 10), leading to roughly similar fixation probabilities on autosomes and on Y chromosomes. Underdominant mutations tend to segregate slightly longer on autosomes on average, especially as more mutations manage to avoid an early purge. However, the mean segregation time remains orders of magnitude smaller than in the overdominant scenarios. Indeed, underdominant mutations are either rapidly purged from the population or, through genetic drift, reach sufficiently high frequencies for homozygotes *aa* to become common, which subsequently facilitates their relatively rapid fixation.

A somewhat opposite pattern emerges in the overdominant scenario (*h* < 0, *s*_0_ < 0 or *h* > 1, *s*_0_ > 0). On the Y chromosome, the mutation maintains a constant selective advantage, whereas on the autosomes its advantage decreases as its frequency increases due to the rising proportion of homozygotes. As discussed above, such conditions generate balancing selection around a metastable frequency on autosomes, in contrast to what is observed on the Y chromosome. Consequently, fixation probabilities are much higher and mean segregation times much shorter on the Y chromosome than on autosomes, and particularly when *h* < 0 and *s*_0_ < 0. Even in cases where the fixation probabilities of overdominant mutations on autosomes and the Y chromosome are similar (*e*.*g*., when *h* = 0.1, *s*_0_ = 100), we can observe some fixations on the Y chromosome and never on the autosomes over the evolutionary timescale considered, due to the large difference in fixation times.

Finally, in the deleterious scenario (*s*_0_ < 0, *h* > 0), which also includes underdominant mutations), fixation probabilities are consistently higher on the Y chromosome, reflecting the stronger genetic drift experienced by Y-linked alleles. Interestingly, however, mean segregation times remain shorter for deleterious mutations on the Y chromosome than on autosomes. This pattern indicates that, even though the few deleterious mutations that do fix on the Y may segregate longer than those which are rapidly purged, they are too rare to increase the mean segregation time above that of autosomal deleterious mutations (see Figure 4**c**).

## 4 Discussion

### The importance of a unified framework for studying the evolutionary trajectories of autosomal and Y-linked mutations

Understanding how mutation fixation probabilities differ between Y-like chromosomes and autosomes is essential for explaining sex chromosome differentiation and its evolutionary consequences, including the build-up of genetic incompatibilities contributing to speciation and the accumulation of sex-linked deleterious mutations underlying many heritable diseases. Y-chromosome specificities in gene transmission, due to their lower effective population size as well as their forced heterozygosity, among other mechanisms, greatly impact the fixation probabilities and mean segregation times of mutations. To compute these key quantities, we used diffusion approximations derived from a biparental two-sex diploid Wright-Fisher model encompassing X and Y sex chromosomes and autosomes. We provided analytical and numerical tools to derive the mutation fixation probability and mean segregation time from this unified framework, for mutations evolving under different kinds of selective regimes.

The fixation probabilities and mean segregation times of mutations depend on the interplay between selection and genetic drift, and their role can be directly established through the coefficients of our diffusion approximations. In the Y chromosome context, no individuals with genotype *aa* are formed and the fate of the mutation only depends on the product *hs*_0_ and on its initial frequency. For example, if *hs*_0_ > 0, selection favours the mutation regardless of its frequency, as the diffusion drift term remains positive for any value of *q*_*t*_ ∈ [0, 1]. Genetic drift counteracts the increase in mutation frequency driven by selection, especially when the mutation is still rare in the population. By contrast, the role of genetic drift in the autosomal context is more complex. While it can also lead to the early loss of the mutation when it fails to reach relatively high frequencies in the first generations, genetic drift can also override selection by driving one allele to fixation even in regimes where selection tends to maintain polymorphism, particularly under strong balancing selection. The framework we provided allows for direct comparisons of fixation probabilities and of mean segregation times between autosomal and Y-linked mutations, as time is scaled identically before taking the large population limit in the two chromosomal contexts. In addition, our framework allows us to calibrate the initial frequencies of autosomal and Y-linked mutations in order to reflect the difference in effective population size of the two chromosome types.

### Recovering classical results in population genetics

Our approach retrieves classical results in population genetics. For instance, deleterious mutations experience higher fixation probabilities on Y chromosomes compared to autosomes, solely due to the increased genetic drift experienced by the Y chromosome. Conversely, we observe that underdominant mutations are more likely to fix on autosomes than on Y chromosomes for a wide range of parameters (*e*.*g. h* ≤ − 1, *s*_0_ ≥ 10). We also show that purely beneficial mutations (*e*.*g. h* = 0.1, *s*_0_ = 100) can reach similar fixation probabilities in both the autosomal and Y-linked contexts (Patwa and Wahl, 2008). Given that mutations arise more frequently on autosomes due to their larger population size, one might expect a higher overall number of fixations on autosomes in this case, although this conjecture must be nuanced in the light of segregation times. Indeed, segregation times are often overlooked, despite their strong influence on the number of fixations actually occurring in a finite time window. In particular, they are not taken into account by the classical substitution rate - yet it is essential for properly interpreting the fixation probability results.

### The importance of the mean segregation time

These time considerations play a particularly important role in the overdominant scenario. Indeed, when *h* > 1 and *s*_0_ > 0, the mutation is still beneficial in the homozygous state but heterozygotes become advantaged comparatively to homozygotes. This has no impact in the Y chromosome context, but greatly modifies the behaviour of the mutation on autosomes, as it becomes disadvantaged once it reaches a high enough frequency to often appear in a homozygous state. This cutoff frequency is a metastable state towards which the process (*p*_*t*_)_*t*≥0_ is attracted, as the associated diffusion drift term is positive when the mutation is rare and negative when it is frequent. Such a metastable state also arises in another overdominant scenario (*h* < 0 *s*_0_ < 0), which explains why we jointly studied these two cases although they looked different at first sight (the mutant homozygotes being fitter than the wild-type homozygotes in the former case and less fit in the latter case). We do observe substantial differences between fixation probabilities in the two cases. These arise from the nature of diffusion processes, which, through rare stochastic excursions, will eventually reach one of the two absorbing states 0 or 1. What actually distinguishes the two scenarios is the range within which the metastable state is found: between 1*/*2 and 1 in the first case (*h* > 1, *s*_0_ > 0) and between 0 and 1*/*2 in the second (*h* < 0, *s*_0_ < 0). Stochastic excursions away from the metastable state, driven by genetic drift, are therefore more likely to bring the mutation to fixation in the first case and to extinction in the second, explaining the discrepancy in fixation probabilities. However, such excursions are unlikely to be observed in a meaningful timescale. Indeed, we observe a very large mean segregation time in both cases (*e*.*g*. an order of 10^69^ generations for *h* = −2 and *s*_0_ = −100). This leads to a rather similar behaviour in the two cases : most mutations are quickly purged from the population, while a small fraction reaches the metastable frequency around which they can oscillate virtually indefinitely. Our framework for deriving the mean segregation time is therefore crucial in such cases for understanding the true evolutionary trajectories of mutations in natural populations.

### Comparison of the Wright-Fisher framework to the individual-based simulations

A consequence of long segregation times is that the theoretical fixation probabilities sometimes deviate from those obtained through individual based-simulations A.1. This is expected, as our simulations are capped at 100,000 generations. Therefore, in cases where the mutation is still segregating at that point, the observed number of actual fixations is 0. Consequently, our theoretical approach can be used in a first instance to determine whether performing time-consuming simulations is worthwhile for a given range of parameters. In addition to scenarios leading to long segregation times, the other situation where our theoretical results differ from simulations is when fixation probabilities are small but not negligible. For example, if the fixation probability is 10^−5^ (which is the case, *e*.*g*., for *N* = 100, *h* = 0.1, *s*_0_ = 100), the probability to actually observe a fixation in 10000 simulations is only 10%, and our simulations indeed ended up without any fixation in this case. This highlights another advantage of our theoretical approach: it enables the estimation of small but non-negligible fixation probabilities in cases where simulations would be prohibitively costly.

### Application to the sheltering effect on Y-like chromosomes

Our analyses show that overdominant mutations are overall more likely to fix in observable time windows on the Y chromosome than on autosomes. This is particularly true in the case where *h* < 0 and *s*_0_ < 0, because the permanent heterozygosity of the Y chromosome prevents the appearance of the disadvantaged homozygotes *aa*. Such protection from homozygous disadvantage by being kept at the heterozygous state due to linkage to a permanently heterozygous locus (or a locus under strong balancing selection) is often called a sheltering effect (Antonovics and Abrams, 2004; Jay et al., 2024). Although single point mutations are unlikely to exhibit true overdominant behavior, both theoretical and empirical studies indicate that large chromosomal rearrangements that suppress recombination are prone to behave as overdominant (Berdan et al., 2023). Indeed, large rearrangements such as inversions tend to capture multiple partially recessive deleterious mutations, which generate a fitness cost when they are homozygous. If these rearrangements confer a fitness advantage when heterozygous due to a lower number of captured recessive deleterious mutations, they can behave as overdominant with resulting negative *h* and *s*_0_. In agreement with previous studies, our results show that this type of chromosomal rearrangements follows distinct evolutionary dynamics on the Y chromosome and on autosomes, with notably higher fixation probabilities on the Y chromosome over observable timescales, which may contribute to the progressive cessation of recombination on the Y chromosome (Hartmann et al., 2021; Tezenas et al., 2023; Jay et al., 2024, 2025).

### Limits and perspectives

Our model focuses on the evolutionary dynamics of one particular mutation (or genomic region) and does not account for the interactions it may have with other parts of the genome. For instance, Y-chromosome degeneration through the accumulation of deleterious mutations involves complex mutation interactions, such as Hill-Robertson interference or genetic hitchhiking (Bachtrog, 2008). Our model also does not take into account possible changes in the mutation fitness (Kimura, 1962; Waxman, 2011), which can be particularly relevant in cases where the mutation segregates for a long time. Other works also tackle the inherent limitations of the Wright-Fisher framework such as the assumptions of a balanced sex-ratio (Shi et al., 2024), a fixed population size (Otto and Whitlock, 1997; Engen et al., 2009) or a large population (Shafiey and Waxman, 2017). Finally, if weak selection can be sufficient to study the deleterious and underdominant cases, we may want to study mutations with a large impact on fitness in the beneficial case, leading to frequencies with non-continuous trajectories (Mavreas et al., 2022).

Nonetheless, our framework with continuous mutation frequency variations could be used to explore several other directions. First, it would be interesting to investigate the effect of varying population sizes on the relative fixation probabilities between the two chromosomal contexts and to look for chromosomal context-dependent effects of population size. For fixed values of *h* and *s*_0_, fixation can be more likely on the Y chromosome in small populations, and more likely on autosomes in large populations, for example. One could also study the fate of mutations that survived early extinction and that are already widespread in the population. It would also be interesting to work with multi-dimensional diffusions in order to study several interacting mutations. Finally, future studies could derive the allele frequency spectrum of each class of mutations on sex chromosomes and autosomes. Comparing these theoretical spectra with empirical observations from Y chromosomes and autosomes may allow the inference of the distribution of dominance coefficients in nature, a long-standing puzzle in evolutionary genetics, that has yet important implications (see the discussion and references in (Mrnjavac et al., 2025)).

## 5 Conclusion

Overall, our study provides a comprehensive and tractable framework for studying the evolutionary fate of mutations on Y chromosomes and for establishing relevant comparisons with the trajectories of autosomal mutations. Beyond simply estimating fixation probabilities, our results illustrate how chromosome-specific features, such as permanent heterozygosity and population size effects, interact to shape the dynamics of mutation segregation and accumulation. Our study underlines the pivotal role of mean segregation time, which can exceed evolutionary timescales in the overdominant scenario, causing mutations to become trapped at intermediate frequencies. This approach offers a versatile tool for improving our understanding of Y chromosome evolution and lays the groundwork for future studies aiming to dissect the complex forces governing genome evolution.

## Fundings and acknowledgments

SB, AO and AV acknowledge partial support from the chaire program “Mathematical modeling and biodiversity” (Ecole Polytechnique, Museum National d’Histoire Naturelle, Veolia Environnement, Fondation X). AO is supported by a grant from Fondation CFM pour la recherche which had no role in the studies or in the decision to publish.

## Code availability

The scripts used to perform individual-based simulations and to compute the fixation probabilities and the mean segregation times are available on GitHub: https://github.com/Arieloffenstadt/Fixation-probabilities-and-mean-segregation-times-on-Y-chromosomes-and-autosomes

## A Annexes

### A.1 Comparison with the results of Charlesworth et al. (1987)

Charlesworth et al. (1987) provided tools to compare the fate of a mutation appearing on an autosome or a sex chromosome. In particular, they compared the autosomal and Y chromosome contexts in the beneficial and deleterious scenarios using two different approximations. In this section, we contrast our results with those obtained in Charlesworth et al. (1987) and discuss the parameter space in which their approximations provide results comparable to ours.

Actually, Charlesworth et al. (1987) studied *substitution rates*, computed as the rate of apparition of a new mutation times the probability that it eventually reaches fixation. These rates are written 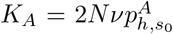 and 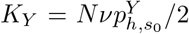 for autosomes and Y chromosomes respectively, with 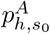 and 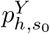 representing the fixation probabilities in the two cases for parameters *h* and *s*_0_. As the rate of apparition of a mutation *ν* is assumed to be the same in all chromosomes, the total rate at which a mutation appears on an autosome (2*Nν*) is thus four times higher than the rate on Y chromosomes (*Nν/*2). The main quantity studied in Charlesworth et al. (1987) is the ratio 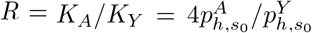. Note also that the selection coefficients they consider are slightly different for Y chromosomes as the heterozygotes have a fitness given by 1 + *s*, instead of 1 + *hs* as in our model. To account for this difference, we will compare the fixation probabilities for given sets of parameters (*h, s*_0_) by fixing *h* = 1 in the Y chromosome context, that is studying the ratio given by 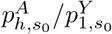.

#### Beneficial mutation

To study the fate of beneficial mutations, Charlesworth et al. (1987) used Haldane’s branching approximation and obtained a ratio of *K*_*A*_*/K*_*Y*_ = 4*h*. In order to compare this result with the ratio we obtain with our approach, we compute the ratio of fixation probabilities on autosomes and Y chromosomes for different values of *s*_0_ and *h* and compare it to the value of *h*. These results, for a population of size *N* = 10, 000, are displayed in Table A.1.

We observe that the theoretical value of *h* is indeed recovered when the product *hs*_0_ is sufficiently large (*hs*_0_ ≥ 10). When *h* and *s*_0_ approach 0, the mutation is almost neutral and the fixation probability is given by the inverse of the population size.

**TABLE A.1:**
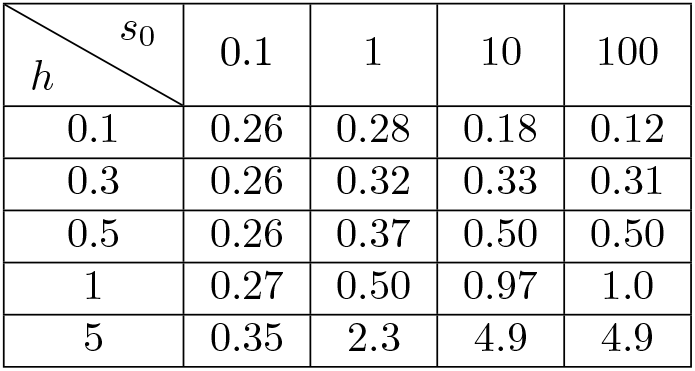
Ratio between the fixation probabilities on autosomes 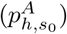 and on the Y chromosome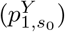 of a single beneficial mutation in a population of size *N* = 10, 000 obtained from our diffusion approximations. Following the results in Charlesworth et al. (1987), we expect a ratio close to *h* on each line.

| $h \backslash s_0$ | 0.1 | 1 | 10 | 100 |
| --- | --- | --- | --- | --- |
| 0.1 | 0.26 | 0.28 | 0.18 | 0.12 |
| 0.3 | 0.26 | 0.32 | 0.33 | 0.31 |
| 0.5 | 0.26 | 0.37 | 0.50 | 0.50 |
| 1 | 0.27 | 0.50 | 0.97 | 1.0 |
| 5 | 0.35 | 2.3 | 4.9 | 4.9 |

**TABLE A.2:** Mean segregation time of a single mutation appearing on an autosome 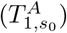 and on a Y chromosome 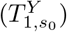 in a population of size *N* = 10, 000, considering different values for *s*_0_ and fixing *h* = 1, obtained from our diffusion approximations.

| $s_0$ | 1 | 10 | 100 |
| --- | --- | --- | --- |
| $T_{1,s_0}^A$ | 0.0020 | 0.0025 | 0.0029 |
| $T_{1,s_0}^Y$ | 0.0026 | 0.0049 | 0.0105 |
| $T_{1,s_0}^A/T_{1,s_0}^Y$ | 1.32 | 1.99 | 3.60 |

This explains the ratio of 0.25 that we observe in the top left part of Table A.1. Our framework therefore allows a continuous exploration of the whole range of parameters, connecting the case of neutral mutations to the case where Haldane’s approximation holds. Similar results are obtained for populations of size *N* = 1000 and *N* = 100, 000.

Furthermore, the classical notion of substitution rate used here does not take into account the time needed by a mutation to actually reach fixation. While the rates of fixation on autosomes and on the Y chromosome may be similar in a given scenario, fixation could happen much quicker on the Y chromosome, thus altering the number of fixation events actually observed within a given time window. Although our framework only allows to study the mean segregation time (*i*.*e*., the mean time until fixation or extinction of the mutation) rather than the mean segregation time conditioned on the mutation eventually fixing, it provides nonetheless a good heuristic indicator.

For example, considering *h* = 1, one should expect that 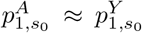 and thus a substitution rate 4 times higher on autosomes than on Y chromosomes. However, we notice in Table A.2 that the mean segregation time is larger on autosomes. In the beneficial mutation scenario and in absence of strong balancing selection (*h* ≤ 1), the mean time to purge is relatively small and of the same order of magnitude in Y chromosomes and autosomes (below fifteen generations). When the probabilities of fixation are close together, the ratio of our mean segregation times is therefore a lower bound for the ratio of the mean times to fixation. In particular, for the set of parameters (*s*_0_ = 100, *h* = 0.1), the ratio is bounded from below by 3.6 (in fact, the simulations presented in Figure A.1 actually ended up with a ratio around 5). This observation reinforces the idea that the mean fixation or segregation times can have an impact on the actual fixation of the mutation on observable timescales.

#### Deleterious mutations

In the deleterious mutation scenario, Charlesworth et al. (1987) used the same diffusion approximation as we do, but performed further Taylor approximations to simplify the expression appearing in (2.12). They obtained a simpler analytical value given by the ratio

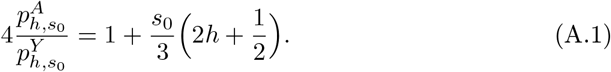

As *s*_0_ < 0, the ratio (A.1) is always below 1, and therefore mildly deleterious mutations are more likely to fix on a Y chromosome than on an autosome. However, we also see that, as the ratio should remain positive, this result applies only to the case where *h* and *s*_0_ are simultaneously relatively small.

Table A.3 compares the formula (A.1) and our results, again for *N* = 10, 000 and different parameter values.

**TABLE A.3:** Comparison between the fixation probability ratio computed in Charlesworth et al. (1987) and that computed with our diffusion approximations, in the deleterious mutation scenario.

| $h \backslash s_0$ | -100 | -10 | -1 | -0.1 |
| --- | --- | --- | --- | --- |
| 0.1 | $3.3e - 44$ | $4.0e - 05$ | 0.17 | 0.24 |
| 0.3 | $2.5e - 44$ | $3.1e - 05$ | 0.15 | 0.24 |
| 0.5 | $1.9e - 44$ | $2.3e - 05$ | 0.13 | 0.24 |
| 1 | $1.5e - 45$ | $5.7e - 06$ | $9.7e - 02$ | 0.23 |
| 5 | $8.9e - 200$ | $3.1e - 21$ | $4.0e - 03$ | 0.17 |

| $h \backslash s_0$ | -100 | -10 | -1 | -0.1 |
| --- | --- | --- | --- | --- |
| 0.1 | $X$ | $X$ | 0.19 | 0.25 |
| 0.3 | $X$ | $X$ | 0.16 | 0.24 |
| 0.5 | $X$ | $X$ | 0.13 | 0.24 |
| 1 | $X$ | $X$ | 0.04 | 0.23 |
| 5 | $X$ | $X$ | $X$ | 0.16 |

(B) Values of the theoretical ratio computed in Charlesworth et al. (1987) and described in (A.1). The $X$ 's represent negative values which can not correspond to a ratio of positive probabilities.

We obtain similar values for when the values of *h* and *s*_0_ are close to 0. When the product *hs*_0_ becomes too large, the analytical formula provided in Charlesworth et al. (1987) becomes negative and cannot be applied. Indeed, when the strength of negative selection increases, the probability of fixation plummets more rapidly to 0 on autosomes than on Y chromosomes. In this case again, our approach allows to study the whole range of parameters from close-to-neutral mutations (ratio close to 1*/*4) to severely deleterious mutations (ratio close to 0).

### A.2 Further simulations

In this section, we present further comparisons between simulations and theoretical results for different parameter regimes, including the four scenarios presented in Table 1. Figure A.1 displays these comparisons, which show overall excellent agreement between our analytical results and simulations, even for small population sizes (from *N* = 100). We observe significant differences in only two situations. First, in some overdominant scenarios (*h* = 10, *s*_0_ = −1; *h* = −1, *s*_0_ = −1), the mean segregation time can be very large on autosomes (see Section 3.2). Since simulations are capped at 100,000 generations, we are unable to observe fixations within this time window. Some mutations are still segregating at the end of the simulations, some of which may be mutations that would eventually reach fixation. Second, when mutations are deleterious (*h* = 0.1, *s*_0_ = −100) or underdominant (*h* = 10, *s*_0_ = −1 or *h* = −10, *s*_0_ = 1), we also observe significant differences, especially for the Y chromosome. This is purely due to statistical sampling, as events with probabilities of 10^−5^ to 10^−7^ are unlikely to be observed in 10,000 trials. This underlines an advantage of our analytical approach, which can provide accurate estimates even for rare events.

**FIGURE A.1:**
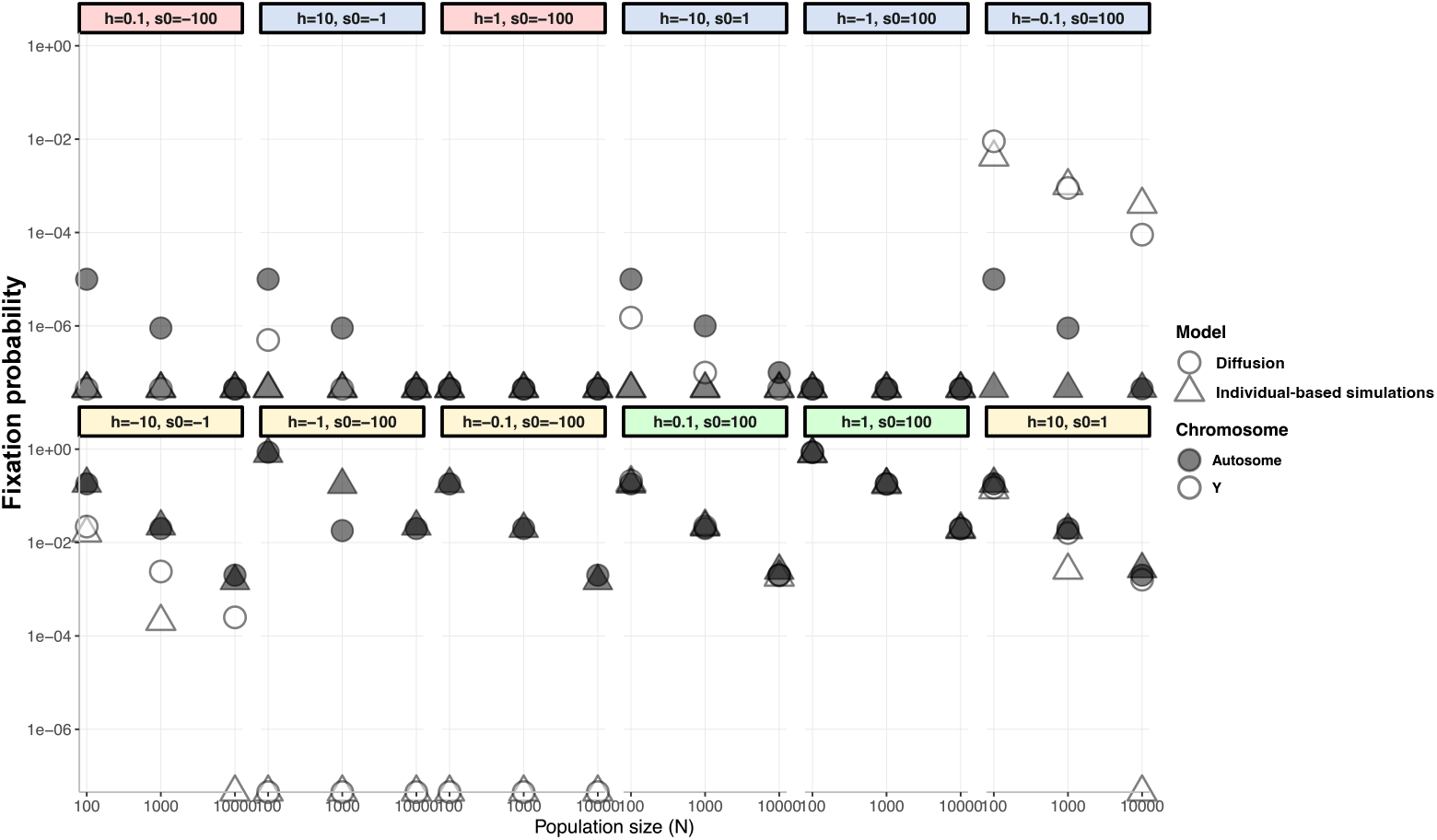
Comparison between the fixation probabilities obtained from the diffusions (*p*_*t*_)_*t*≥0_ (2.10) and (*q*_*t*_)_*t*≥0_ (2.11) and the results of 100,000 individual-based simulations described in Section 2.5. We study three population sizes *N* = 100, *N* = 1000 and and *N* = 10000, for which each set of parameter (*h,s*_0_) induces differences in fitness given by 1 for the genotype *AA*, 1 + *hs*_0_*/N* for the genotype *Aa* and 1 + *s*_0_*/N* for the genotype *aa*, in accordance with (2.1). For each *N*, the simulations start with a single mutation in a diploid population of size *N*, and the diffusions start at initial frequencies *p*_0_ = 1*/*(2*N* ) and *q*_0_ = 2*/N* . White (*resp*., gray) triangles represent the fixation probabilities obtained with the simulations for a mutation appearing on an autosome (*resp*., on a Y chromosome), while white (*resp*., gray) circles represent the fixation probabilities computed with the diffusion approximations for a mutatiton appearing on an autosome (*resp*., on a Y chromosome). The parameter sets studied include the four scenarios described in Table 1, which are made explicit by the coloured backgrounds (green for beneficial, red for deleterious, yellow for over-dominant and blue for underdominant).

### A.3 Proofs

In this section, we provide details on the strategy of the proof of the diffusion approximations obtained in Section 2.2 and 2.3. The key tool we use is the Stroock-Varadhan Theorem (see, *e*.*g*., Theorem 7.1. in Chapter 8 of (Durrett, 2008)). In a second time, we compute the fixation probabilities and mean segregation times of these diffusions.

#### A.1.3 Convergence to diffusions

In Section 2.2, we encoded the frequencies of a mutation in a population of *N* diploid individuals by the Markov chains 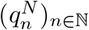 and 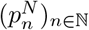, respectively in the Y chromosome and autosomal contexts, and we described their dynamics.

As for large *N* values, the allele frequency change between two generations has a variance of order 𝒪 (1*/N)*, we use the classical strategy of accelerating time and work with the rescaled processes 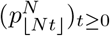 and 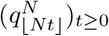 defined in Section 2.3. They are piecewise constant continuous-time processes evolving in a sped-up timescale where 1 unit of time corresponds to *N* generations.

Our goal is to show that, as 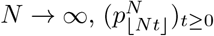 and 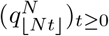 converge towards the diffusion processes described in (2.10) and (2.11).

To this end, we apply Theorem 7.1 in (Durrett, 1996, chap. 8), which helps compute the coefficients of the limiting diffusions through the one-step changes in frequencies that we already derived in Section 2.2. Since 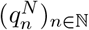 and 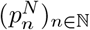 are time-homogeneous, it is actually enough to compute the changes in expectation and variance of the frequencies in the first generation, conditionally on the initial frequency.

More precisely, the conditional expectation of the variation of the frequency from generation 0 to generation 1, multiplied by a factor *N* → ∞, converges as *N* to the drift coefficient of the limiting diffusion. Similarly, the conditional variance of the variation of the frequency from generation 0 to generation 1, multiplied by a factor *N* → ∞, converges as *N* to the variance coefficient of the limiting diffusion. We therefore compute these quantities.

##### The Y chromosome case

Denote, for each *N* ∈ ℕ, *β*^*N*^ = *hs*^*N*^ = *hs*_0_*/N* and 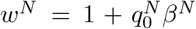. In the Y chromosome context, we have seen in (2.9) that conditionally on 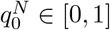, we have 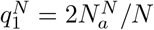 with

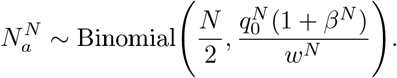

As

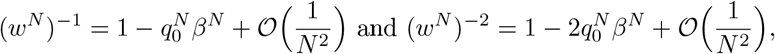

we have that

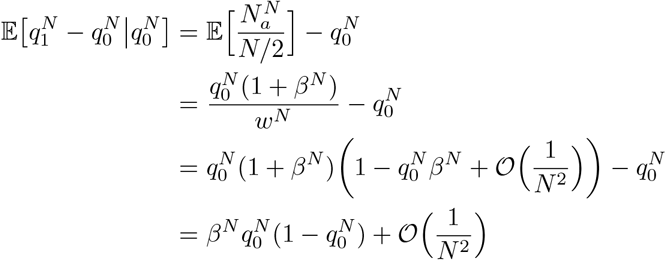

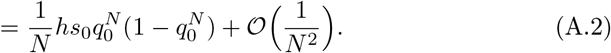

Similarly we obtain that:

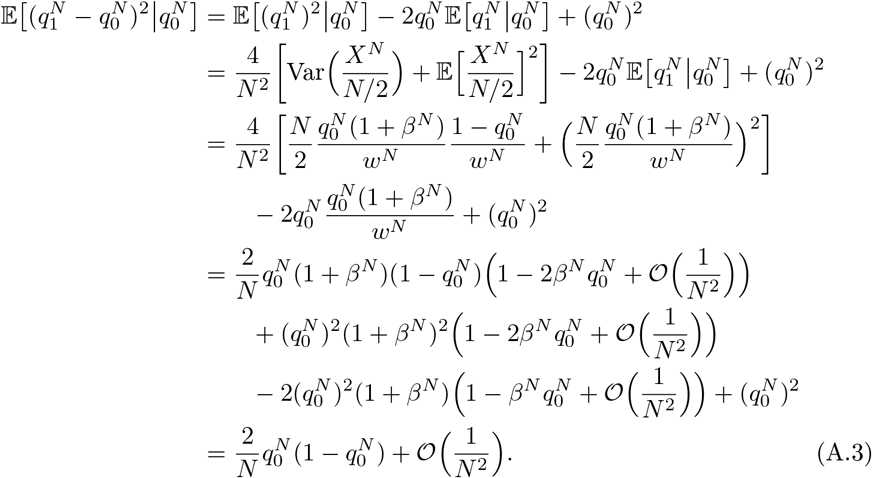

Multiplying the expressions in (A.2) and (A.3) by *N* and letting *N* tend to infinity, the terms of order 𝒪(1*/N)*^2^ vanish and we obtain respectively the drift and variance coefficients of the diffusion defined in (2.11).

##### Autosomes

Because it simplifies the computations, we consider the Markov chain 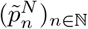 _N_ describing the frequency of the ancestral allele *A* and compute the coefficients of the associated diffusion 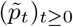. Similarly to the expression (2.6), we have that conditionally on its initial frequency 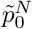,

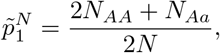

Where

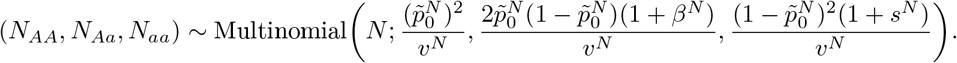

In the previous expression,

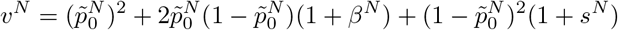

can be written *v*^*N*^ = 1 + *s*^*N*^ *K* with 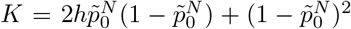 being of order 𝒪 (1). Following the same steps as in the Y chromosome context, we compute that

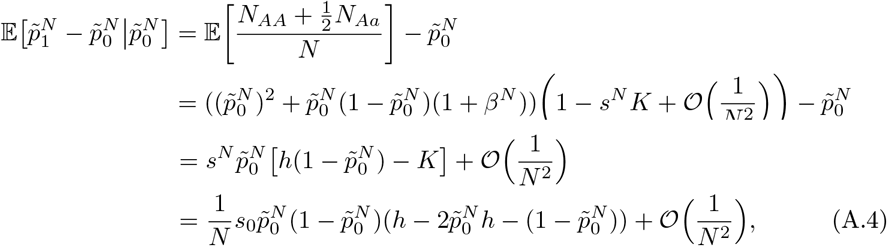

and that

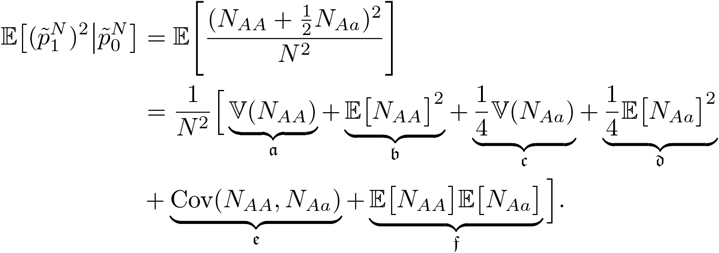

Computing each term separately, we have that

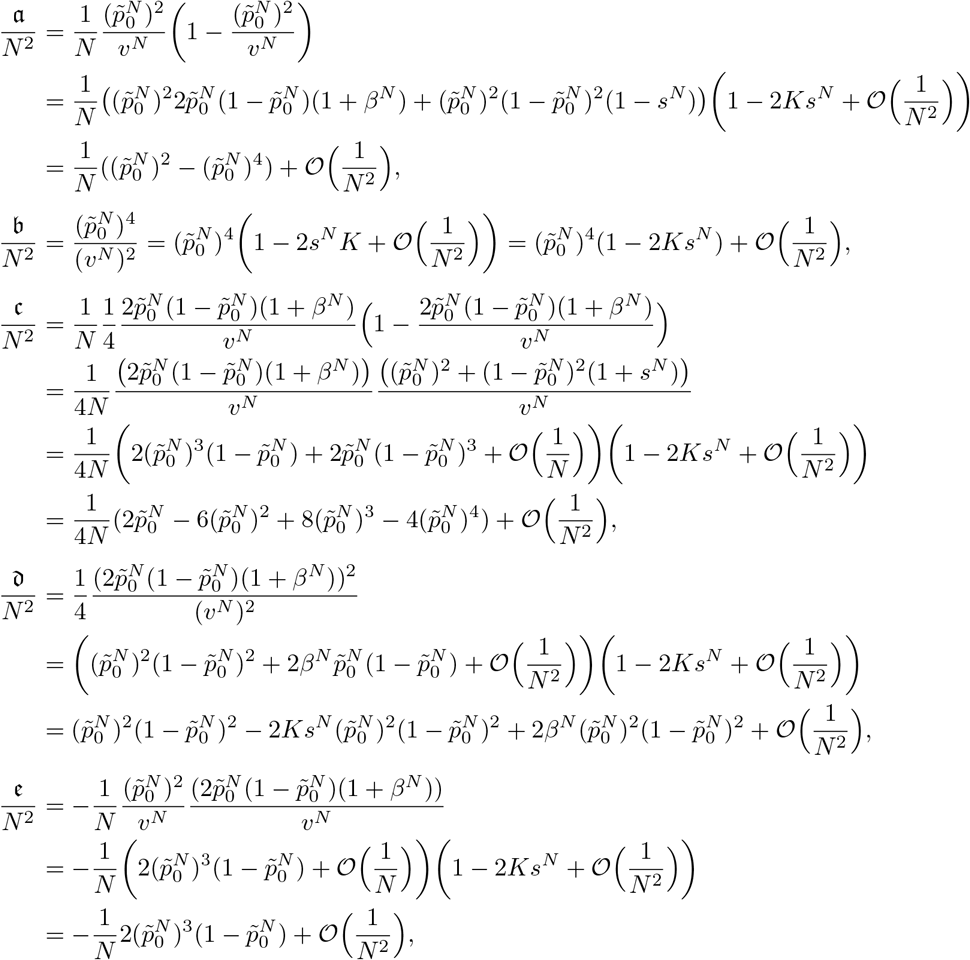

and finally

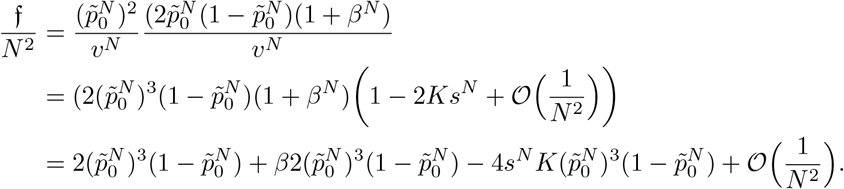

Using (A.4), we also have that

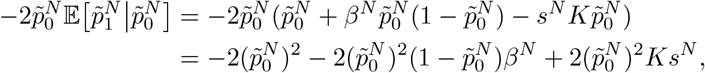

and we finally obtain

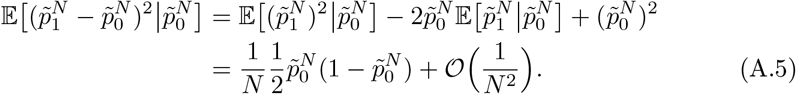

Once again, multiplying both results (A.4) and (A.5) by *N*, we obtain that the diffusion 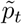 describing the frequency of the ancestral allele *A* is given by

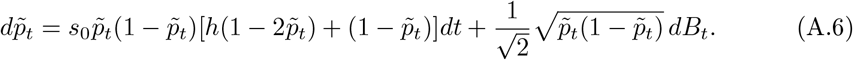

Using that 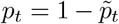 at any given time *t* and plugging this relation in (A.6) (noting that 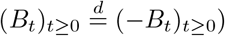, we recover for (*p*_*t*_)_*t*≥0_ the equation (2.10).

#### A.3.2 Fixation probabilities and mean segregation times

Now that we have derived diffusion approximations for the trajectory of mutation frequencies in our Wright-Fisher model, in the Y chromosome and autosomes contexts, we apply classical results on one-dimensional diffusions to derive two key quantities. First, we compute the fixation probability as the probability that the diffusion reaches the absorbing state 1 before being absorbed at 0. Second, we compute the mean segregation time as the average time the diffusion spends away from the absorbing states 0 and 1. To do so, we follow the approach described in Section 3.3 in Etheridge (2011), and compute the *scale functions S*_*p*_ and *S*_*q*_ for the diffusions. We obtain, for every *x* ∈ (0, 1),

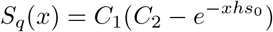

and

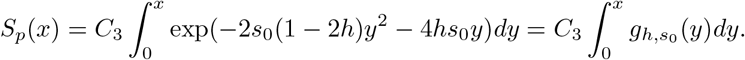

The constants *C*_1_, *C*_2_ and *C*_3_ depend explicitly on the parameters *h* and *s*_0_. As they cancel out in the formulae below, we do not make their values explicit. Lemma 3.14 in Etheridge (2011) then allows us to compute the following fixation probability.

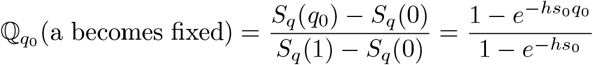

which yields the expression (2.13). This result was expected in the light of the classical Wright-Fisher model with selection. Similarly, we have that

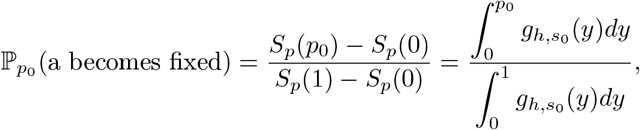

which indeed corresponds to (2.12).

Define now 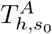 (*resp*.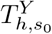) as the expectation of the first time at which *p*_*t*_ (*resp. q*_*t*_) reaches the absorbing state 0 or 1, when the selection parameters are given by *h* and *s*_0_ and the initial frequency by *p*_0_ (*resp. q*_0_). Applying Theorem 3.19 in (Etheridge, 2011, p.44) with the test function *g*(*x*) ≡1, we obtain that the mean segregation time in the Y chromosome context is given by

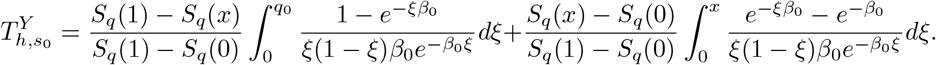

Writing

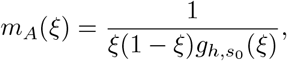

we obtain the following expression for the mean segregation time in the autosomal context:

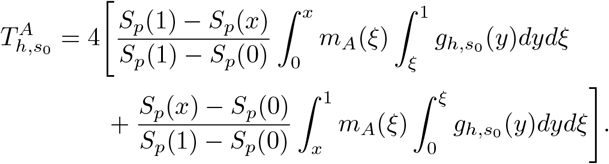

